# Unraveling molecular mechanisms for specification of blastomere identity and neural fate in the annelid *Capitella teleta*

**DOI:** 10.64898/2026.11.13.766470

**Authors:** Davila-Sandoval. Johnny A, Javier Tabima, Néva P Meyer

**Affiliations:** Biology Department, Clark University, 950 Main Street, Worcester, MA 01610; Department of Evolutionary Developmental Genetics, Georg-August-Universität Göttingen, Justus-von-Liebig-Weg 11, Göttingen, Germany 37077

**Keywords:** neural specification, spiral cleavage, blastomere identity, *Capitella teleta*, Annelida, Spiralia

## Abstract

The evolution of central nervous systems remains highly debated as we incorporate data on developmental mechanisms in understudied taxa. Yet how this process occurs in the third major bilaterian clade called Spiralia (e.g., annelids, mollusks) remains inconclusive. Using blastomere isolations, we previously found that in the annelids *Capitella teleta* and *Platynereis dumerilii*, ectodermal blastomeres fated to generate the brain and ventral nerve cord can form neural tissue after being isolated. However, in *C. teleta,* nerve cord formation can be inhibited or rescued by other blastomeres. To ascertain the molecular mechanisms underlying this process, we performed RNAseq on isolated blastomeres in *C. teleta* at the 8-, 16- and 32-cell stages and found differential expression of components of canonical Wnt signaling across the animal-vegetal axis when conditional neural specification is occurring. Furthermore, asymmetrical localization of *TALE* and *FOX* transcripts along the embryonic axis suggests a role in establishing blastomere identity, as observed in mollusks. Our data suggest the integration of an ancestral gene regulatory network of *FOXQ2-*cWnt and lineage-specific genes during early development, linked to the emergence of spiral cleavage.

**Summary statement:** Asymmetrical expression of Spiralia-specific transcription factors and FOXQ2-cWnt dynamics along the embryonic axis in *Capitella teleta* unveil an integration of ancestral and lineage-specific molecular mechanisms underlying blastomere identity in spiralians.

## Introduction

Spiralia is a major clade within bilaterians that displays a wide array of body plans, but underlying molecular mechanisms controlling development of this astounding diversity are still largely unknown. In addition to molecular evidence supporting the existence of this clade (Giribet, 2008; Marlétaz et al., 2019), many taxa share an ancestral mode of early development called spiral cleavage. Across annelids, mollusks, and other spiralians, stereotypical divisions lead to similar blastomere arrangements and lineages (Ackermann et al., 2005; Hejnol et al., 2007; Meyer et al., 2010; Render, 1997); however, the molecular underpinnings of this process are still poorly understood. During spiral cleavage, D-quadrant cells including the organizer are specified either conditionally or autonomously. In the former, inductive interactions among neighboring blastomeres are necessary to specify the D-quadrant. In the latter, asymmetric inheritance of maternal determinants specifies the D-quadrant (Amiel et al., 2013; Lambert et al., 2016; Lambert & Nagy, 2001, 2001; Lanza & Seaver, 2020b, 2020a). The conservation of cleavage patterns cross-phyla and evolutionary shifts between different modes of specification make spiral cleavage an ideal system to study how conserved cell lineage programs can lead to different body plans in metazoans.

Spiral cleavage is a pattern of division characterized by alternating 45° shifts in the mitotic spindle with respect to the embryonic axis leading to a spiral-like arrangement of cells in the animal pole (Fig. 1A) (Martín-Durán & Marlétaz, 2020). During this process, four quadrants of the embryo are formed (A–D) at the 4-cell stage. In the subsequent division to the 8-cell stage, one daughter cell is localized to the animal pole and tends to be smaller (micromeres, 1a–1d), and the other daughter cell is larger and localized to the vegetal pole (macromeres, 1A–1D). As blastomeres continue to divide, their identity is assigned based on their lineage (e.g., 2q = second-quartet micromeres = 2a, 2b, 2c, and 2d, see Fig. 1B). Conservation of spiral cleavage across multiple taxa allows for the identification and fate mapping of homologous blastomeres in animals with unequal cleavage, which can be used to understand evolution of cell fate specification.

**Figure 1.**
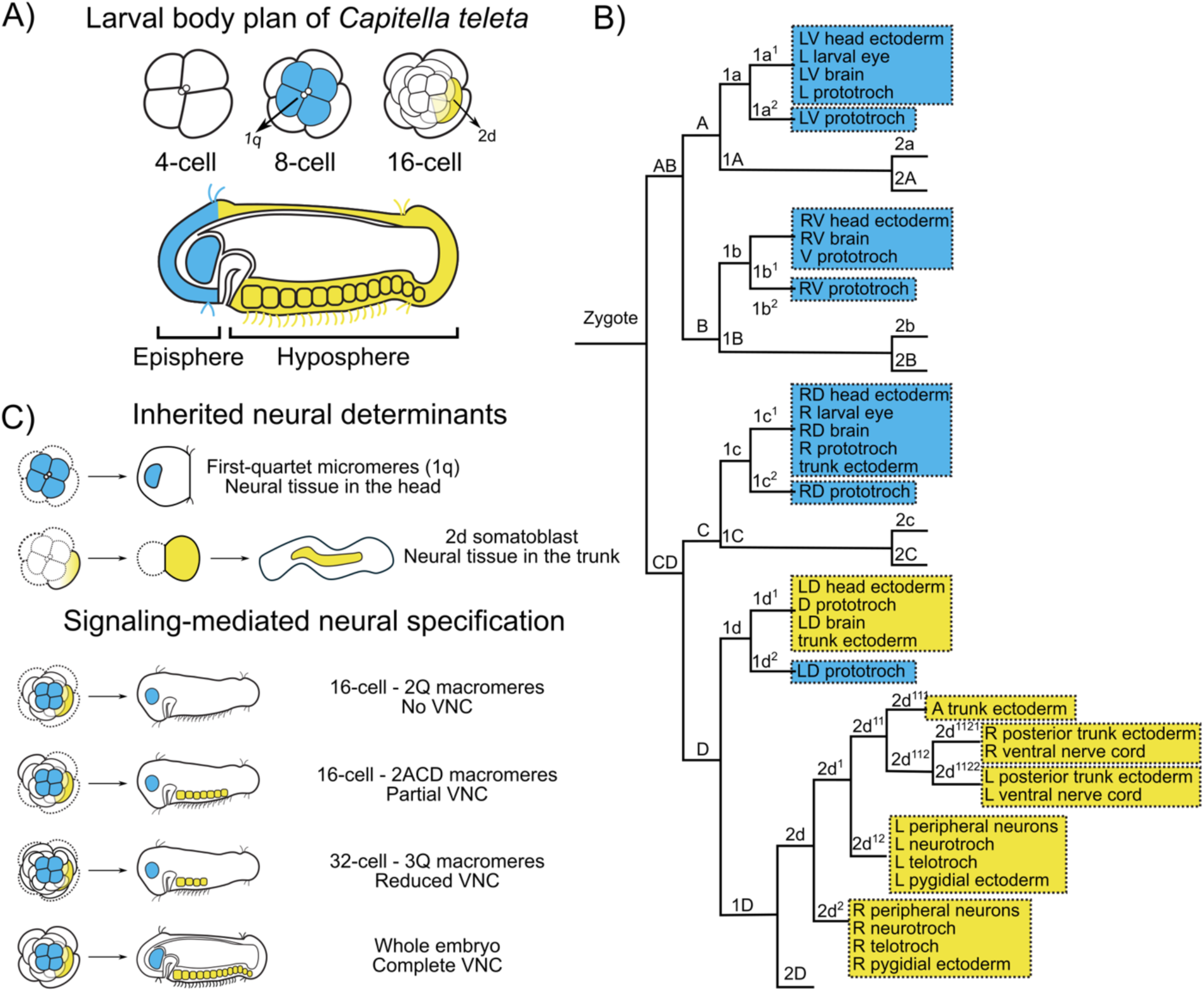
BMP signaling and CNS specification in the annelid *Capitella teleta*. A) Larval body plan of *C. teleta* based on fate-mapping. Top: Early embryonic stages indicating the 1q micromeres (blue) and the 2d somatoblast (yellow). B) Cell lineage diagram for *C. teleta* highlighting the fates of the 1q (blue) and 2d (yellow) blastomeres. C) Isolation experiments performed in *C. teleta* to test for inherited vs. signaling-mediated factors. Top: 1q and 2d micromeres form neural tissue in isolation. Bottom: Specification of VNC by 2d is repressed by other 2q animal micromeres and restored by addition of any one 2Q macromere (e.g., 2q + 2B). Isolating the micromeres including 2d from the macromeres at the 32-cell stage results in formation of a reduced VNC.

In Spiralia, fate specification appears to occur during early embryogenesis, as isolated blastomeres continue to make their fated tissues in the absence of outside signals (Carrillo-Baltodano & Meyer, 2017; Clement, 1962; Penners, 1924; Webster et al., 2026; E. B. Wilson, 1904). Furthermore, certain tissues and cells seem to come from similar blastomeres across multiple taxa. For instance, the mesoderm has a dual origin, ectomesoderm and endomesoderm. In multiple species, including the nemertean *Cerebratulus lacteus* (Henry & Martindale, 1998), the polyclad *Hoploplana inquilina* (Boyer et al., 1996, 1998), the mollusk *Crepidula fornicata* (Hejnol et al., 2007), and the annelid *Platynereis dumerilii* (Ackermann et al., 2005), endomesoderm originates from 4d, known as the mesentoblast. In annelids, including the unequal cleavers *P. dumerilii* and *Capitella teleta*, the brain generally originates from 1q micromeres while the ventral nerve cord (VNC) is formed by micromere 2d (Ackermann et al., 2005; Meyer et al., 2010) (Fig. 1A, B). These similarities provide a basis to incorporate molecular evidence to test lineage homology cross-species and even cross-phyla.

During embryonic development, some blastomere(s) also influence specification of neighboring cells. For example, the identity and timing of the organizer can vary across taxa, but it is generally restricted to one or two blastomeres in the D quadrant at the 16- to 64-cell stage. While its D-quadrant origin is consistent, the mechanism of specification varies between equal and unequal cleavers. In equal-cleavers, such as the annelid *Owenia fusiformis* and *Spirobranchus lamarcki,* quadrants are morphologically indistinguishable until conditional signaling establishes one 4q micromere as the 4d organizer, resulting in ERK1/2 activation. 4d then patterns dorsal-ventral (D-V) fates via BMP signaling (Carrillo-Baltodano et al., 2026; Seudre, Carrillo-Baltodano, et al., 2022). In unequal-cleavers such as the mollusks *I. obsoleta* and *Crepidula fornicata*, specification of the organizer still occurs in a D-quadrant cell and involves ERK1/2 activation, but the process relies on maternally-inherited factors and D-quadrant cells are visibly larger prior to ERK1/2 activation (Amiel et al., 2013; Carrillo-Baltodano et al., 2026; Henry & Perry, 2008; Lambert & Nagy, 2001). This indicates that specification occurs during early development in Spiralia with some major features conserved cross-phyla and with some differences depending on the type of spiral cleavage.

Neural specification has been widely studied in a subset of species in vertebrates and insects as they generally show similarities at a mechanistic level. In many vertebrates, central nervous system (CNS) specification follows the “default” model, where previous signaling events influence ectodermal cells to differentiate into neural tissue unless they receive inhibitory signals (Stern, 2005). In this model, a combination of FGF activation and BMP and Wnt inhibition are crucial for CNS specification as part ofD-V axis establishment (Alvarez et al., 1998; Bertrand et al., 2003; Delaune et al., 2005; Slack, 2024; S. Wilson et al., 2001). In insects, BMP/Dpp diffusion along with a counter gradient of Chordin/Sog and Tsg and activation of EGF signaling is required for D-V patterning and the establishment and maintenance of the neurogenic domain (Bressan & Araujo, 2022; Gómez-Skarmeta et al., 2003; Lynch & Roth, 2011; Mizutani et al., 2006; Von Ohlen & Doe, 2000). Interestingly, spiralians display some major divergences in the interplay of signaling pathways during neural specification.

In Spiralia, BMP does not seem to have a conserved role as an anti-neural signal, and D-V axis formation is likely uncoupled from CNS specification, which has been discussed previously (Knabl et al., 2026; Webster et al., 2021; Webster & Meyer, 2024). In annelids, there is no functional evidence supporting a function of BMP signaling in repressing neural fate. For instance, in the annelid *Owenia fusiformis*, BMP signaling instead specifies dorsal-posterior fates in larvae (Carrillo-Baltodano et al., 2026). In the annelid *C. teleta*, using blastomere isolations, we previously found that isolated individual 1q micromeres (8-cell stage) or 2d (16-cell stage) form neural tissue (Fig. 1C) (Webster et al., 2026). However, VNC fate in *C. teleta* is affected by external signaling as isolation of all micromeres including 2d at the 16-cell stage (−2Q) results in larvae with a brain but no VNC while isolation of all micromeres plus any one macromere rescues VNC formation (Fig. 1C) (Carrillo-Baltodano & Meyer, 2017). Furthermore, treatment with rBMP4 after birth of 4d led to the formation of an ectopic eye and brain lobe from micromere 1b, while treatment at the 16-cell stage resulted in a radialized episphere and brain (Webster et al., 2021). Blastomere isolation experiments had similar results in *Platynereis dumerilii,* suggesting conserved mechanisms in neural fate specification (Webster et al., 2026). Furthermore, manipulation of BMP signaling in this species during cleavage stages showed expansion of dorsal or ventral fates only in the episphere (Carrillo-Baltodano et al., 2026). In other spiralians, the effect of BMP on D-V axis formation is generally conserved but its role in neural fate specification is unclear (Lambert et al., 2016; Lyons et al., 2020; Tan et al., 2022). Based on this data, it is possible that a different molecular mechanism has evolved to specify neural fate, one that is uncoupled from D-V axis specification and does not rely on BMP inhibition.

To better understand the molecular mechanisms controlling blastomere identity, organizing activity, and neural specification in spiralians, we examined gene expression dynamics during early development of *Capitella teleta.* The wide array of resources available for this species make it amenable for the study of cell fate specification and CNS evolution (Carrillo-Baltodano & Meyer, 2017; Davies et al., 2026; Meyer et al., 2010; Meyer & Seaver, 2010). Based on our previous data, we generated a model for the specification of the CNS in *C. teleta* (Fig. 1C). With this as baseline, we focused on identifying the transcription factors and signaling pathways involved in this process. To achieve this, we generated transcriptomic profiles of the cells that originate the brain (1q) and nerve cord (2d) via RNA-sequencing (RNA-seq). Furthermore, we generated RNA-seq data in combinations of blastomeres where the formation of the nerve cord is either rescued, repressed, or formed normally, and we identified transcription factors and pathway components that are potentially involved. Our data show the presence of transcription factors (*FOX* and *TALE*), signaling pathway components (cWnt, TGF-β), and a gene hierarchy that has been observed in other bilaterians. This suggests a combination of an ancestral GRN involving *FOXQ2-*cWnt and the integration of lineage-specific mechanisms to specify the CNS and establish blastomere identity in the annelid C. *teleta.* Furthermore, it provides important insights into the specification of blastomere identity during spiral cleavage, mRNA segregation, and organizer activity, which are central to understand the origin of body plans in Spiralia.

## Results

### Transcriptomic profile of first-quartet (1q) micromeres, which generate brain tissue in isolation

We previously demonstrated that all first-quartet micromeres (1q) or individual 1q micromeres 1a, 1b, 1c, or 1d can form neural tissue in isolation (Fig. 2A) (Carrillo-Baltodano & Meyer, 2017; Webster et al., 2026). To identify the factors that are inherited in these micromeres, we generated a transcriptomic profile for isolated 1q and 8-cell embryos and performed pairwise comparisons. We used a similar nomenclature for isolations as in Webster et al., (2026), with some modifications (Table S1). Previous data in *C. teleta* suggested that the maternal-to-zygotic transition (MZT) in this species is between the 8- and 16-cell stage based on a sharp increase in gene expression at the 16-cell stage (Liang et al., 2025). In our analysis, we found a limited number of differentially-enriched genes (DEGs) across both treatments. We found 14 genes to be enriched in 1q including *Ct-zic*, *Ct-foxQ2A, Ct-foxQ2B* and *Ct-fox-orphan*, while 38 genes were enriched in the whole-embryo including several members of the Forkhead box family of transcription factors (*FOX*) (*Ct-foxE, Ct-foxL2/3*) and the three amino-acid loop extension homeobox class of transcription factors *(TALE)* (e. g., *Ct-taleVIIIA1, Ct-taleVIIIB2, Ct-taleVIIIB3*) (Fig. 2B, Table S2). From our list of DEGs, we performed a gene ontology enrichment analysis and found that in enriched transcripts in 1q, four gene sets were over-represented including those for cell fate commitment (Fig. 2C). To explore potential regulatory factors that might act as maternal determinants along the animal-vegetal axis, we extracted a region of 2750 bp flanking the transcription start site (2500 bp upstream and 250 bp downstream) for all transcripts enriched in 1q vs 8-cell and performed motif enrichment analysis to identify binding sites using the Metazoan JASPAR database (Bailey et al., 2015). Using FIMO, we found 66 binding sites for SMAD factors in most TALE and FOX genes (Table S3). Using MEME, we found a motif with a C2H2 zinc-finger transcription factor binding motif near both 1q-enriched genes (14/14, e-value = 1.75E-003) (Fig. S1) and 8-cell-enriched genes (27/38. e-value = 1.58E-002) (Fig. S2). Although we did not identify BMP signaling pathway components besides *Ct-nogginA* in 8-cell embryos, the presence of SMAD binding sites suggests a possible role of BMP signaling in quadrant specification as shown via BMP protein incubation experiments (Webster et al., 2021). All in all, this suggests that brain specification relies on inherited factors during spiral cleavage including *FOX* and *TALE* mRNAs.

**Figure 2.**
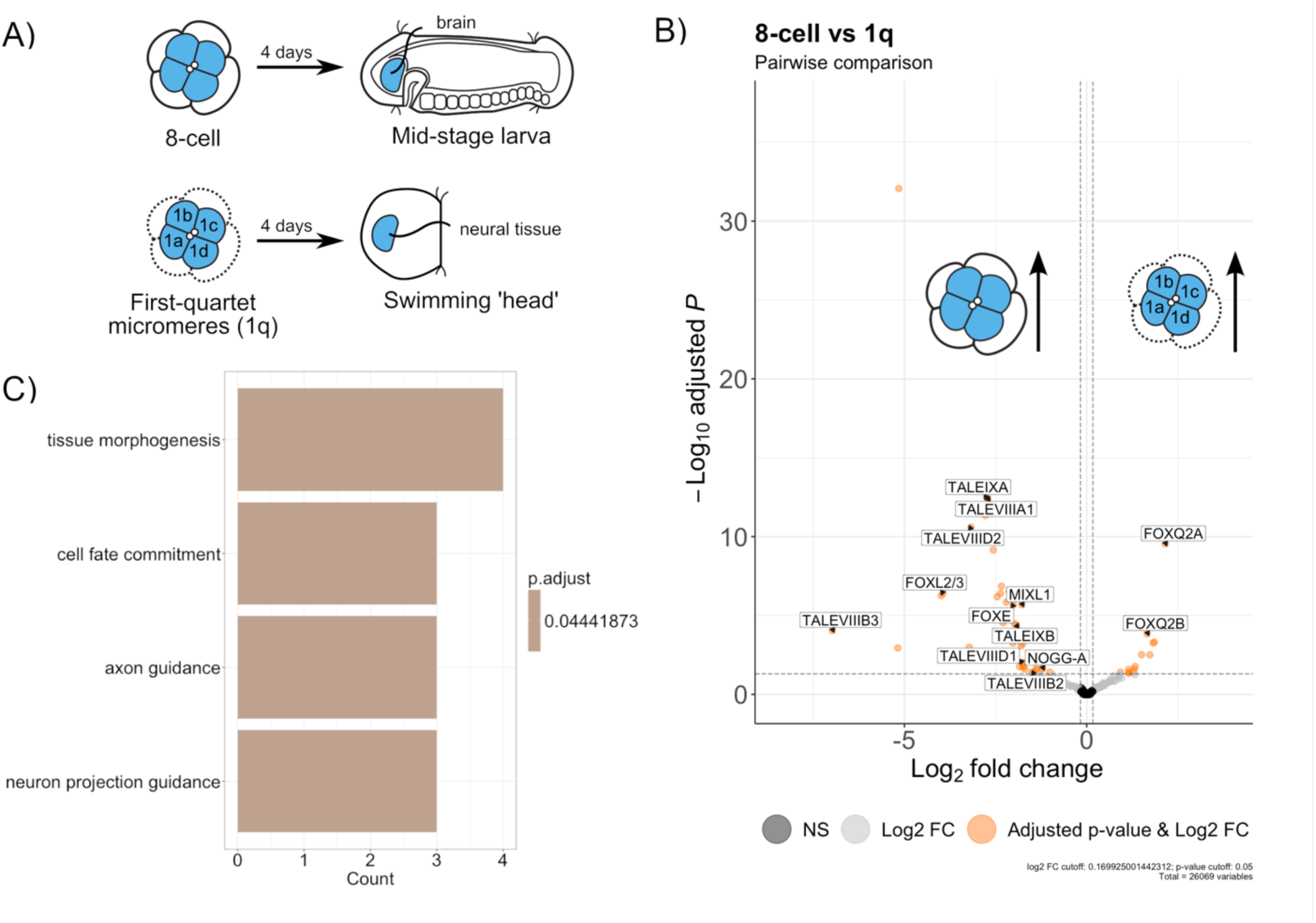
*TALE* and *FOX* genes are differentially enriched across the animal-vegetal axis at the 8-cell stage in *Capitella teleta*. A) Experimental design for transcriptomic analysis. Top: 8-cell embryo, with first-quartet micromeres (1q) labeled in blue (left), forming the brain in a stage 6 larva (right). Bottom: Isolated 1q micromeres with macromeres removed (dotted lines) forming a swimming head with neural tissue (right, blue). B) Volcano-plot showing down-(left) and up-regulated (right) genes in 1q, showing mainly *FOX* and *TALE* genes differentially-enriched. Thresholds for significant differential expression are: Log2 FC > 0.125, adjusted p-value < 0.05. Legend: non-significant differential expression (black), adjusted p-value > 0.05 (gray), significant differential expression (orange). C) Bar plot showing significantly enriched gene ontology terms (adjusted p-value < 0.05) in genes up-regulated in 1q micromeres. Adjusted p-value per ontology term is indicated on the legend in the right.

### Transcriptomic profile of micromere 2d, which is the D-V organizer and forms the VNC in isolation

Signaling centers are present across metazoans and are responsible for fate specification, morphogenesis and formation of the axes. In *C. teleta*, micromere 2d has been identified as both the primary somatoblast and an “organizing cell” (Amiel et al., 2013). We previously showed that 2d can form neural tissue after being isolated from the other blastomeres (Webster et al., 2026). To identify what is inherited in this cell and how it compares to the entire embryo, we generated a transcriptomic profile for 2d. To isolate 2d, the mother cell, macromere 1D, was first isolated from 8-cell embryos. Once this blastomere divided into macromere 2D and micromere 2d, we removed 2D, which was larger and contained yolk granules (Fig. 3A). In parallel, 16-cell embryos from the same brood were also collected. After samples were sequenced, we performed a pairwise comparison of 2d vs 16-cell embryos for differential expression analysis. We found a total of 391 transcripts enriched in 2d and 276 enriched in 16-cell embryos (Fig. 3B). We observed multiple transcription factors enriched in 2d, including *FOX* (*Ct-foxL2/3, Ct-foxK, Ct-foxq2C, Ct-foxq2D, Ct-fox-orphan*) and *TALE* genes (*Ct-taleIV-like, Ct-taleVIIIB2*). Interestingly, from all up-regulated genes in 2d, *Ct-foxq2D* transcripts also were found to be present at high levels in oocytes, zygotes, and early cleavage stages, suggesting a role as a maternal determinant (Fig. S3). We also found transcription factors that were enriched in whole 16-cell embryos, including *TALE* (e. g., *Ct-taleXIIA1, Ct-taleXIIA2, Ct-taleXIIB1*), *FOX (Ct-foxP1, Ct-foxH, Ct-fox-orphan)* and *Ct-gataB1*. Expression of the *Ct-gataB1* in 2Q macromeres at the 16-cell stage and subsequently in the endoderm was previous published (Boyle & Seaver, 2008), validating differential expression of this gene. In our binding site analysis, we found multiple enriched motifs with binding sites for different classes of transcription factors. In transcripts enriched in 2d, we found motifs including binding sites for transcription factors with an NKX (201/391, e-value = 2.0E-004) or a CREB DNA binding domain (114/391, e-value = 2.0E-004). Both classes of transcription factors are expressed during early development (Fig. 3C). Altogether, this indicates that 2d expresses several FOX and TALE transcription factors with NKX homeodomain and CREB transcription factors as potential master regulators. As observed for 1q, a discrete set of transcription factors are enriched in 2d, suggesting that individual blastomeres have unique molecular identities during unequal spiral cleavage.

**Figure 3.**
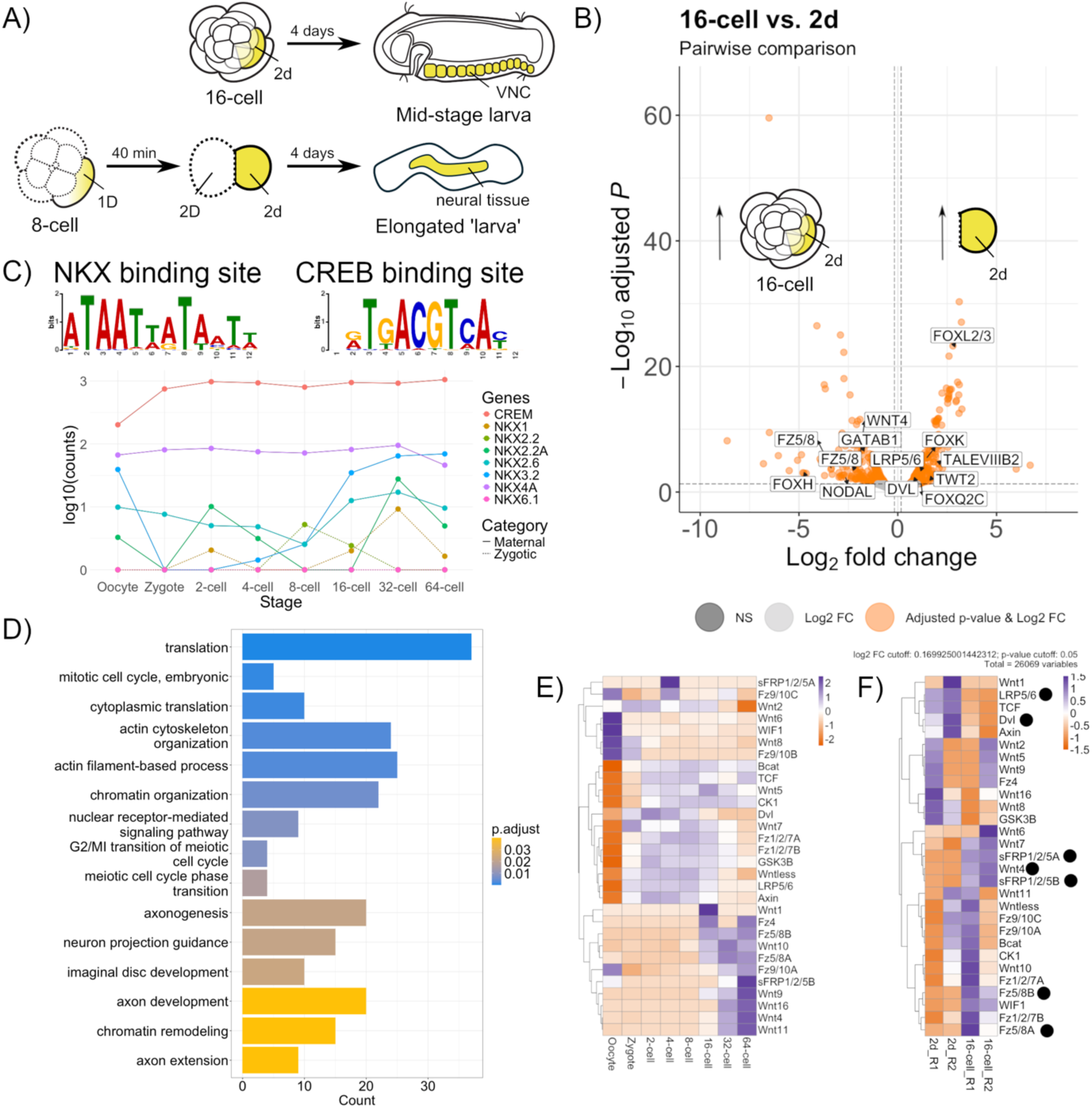
A unique set of *TALE* and *FOX* transcription factors and different canonical Wnt signaling pathway components are enriched in the 2d somatoblast, A) Experimental design for transcriptomic analysis. Top: 16-cell embryo, with the somatoblast 2d labeled in yellow (left), forming the VNC in a stage 6 larva after ∼4 days (right). Bottom: Sequence of isolations to obtain 2d. First, macromere 1D was isolated by removing all the remaining blastomeres (dotted lines), then micromere 2d was isolated by removing 2D (dotted lines). This cell gives rise to trunk ectoderm including neural tissue. B) Volcano-plot showing down-(left) and up-regulated (right) genes in 2d, showing again *FOX* and *TALE* genes differentially-enriched. Thresholds for significant differential expression and legends are as in Figure 2B. C) Up: Consensus binding sites for NK and CREB found in motifs enriched in up-regulated transcripts in 2d. Down: Transcriptomic expression during early development of candidates with aforementioned domains. D) Bar plot showing the top 15 significantly enriched gene ontology terms (adjusted p-value < 0.05) in genes up-regulated in the 2d somatoblast. Adjusted p-value per ontology term is indicated on the legend in the right (orange to blue). E) Heatmap for z-scores of average normalized counts of canonical Wnt signaling (cWnt) pathway components during spiral cleavage (Liang et al., 2025). F) Heatmap for z-scores of average normalized counts of cWnt components in all samples used for 2d vs 16-cell pairwise comparison. Black circles: differentially-enriched in our study.

To understand how 2d might to signal other cells, we also identified signaling pathway components in the list of differentially-enriched genes (DEGs). Using a functional enrichment analysis, we found gene sets related to chromatin remodeling and organization in 2d-enriched transcripts (Fig. 3D). Furthermore, we found differential enrichment of several components of the canonical Wnt signaling pathway (cWnt) in 2d and 16-cell embryos (Fig. 3B). To explore this further, we annotated cWnt pathway components and identified their levels of expression in 2d (our data) and in whole embryos throughout spiral cleavage using the developmental transcriptome available for *C. teleta* (Liang et al., 2025). In 2d, we observed enrichment of *Ct-dishevelled* and *Ct-lrp5/6*, while in the whole-embryo, we observed an enrichment of *Ct-wnt4, Ct-sfrp1/2/5A, Ct-sfrp1/2/5B, Ct-fz5/8A* and *Ct-fz5/8B*: all members of the canonical Wnt signaling pathway (cWnt) (Fig. 3B). Based on transcriptomic data across spiral cleavage in *C. teleta* (Liang et al., 2025), we identified specific combinations of ligands and receptors with similar temporal patterns of expression (Fig. 3E), including *Ct-wnt4* and *Ct-sfrp1/2/5B*, which were found to be enriched in whole-embryos (Fig. 3B). During spiral cleavage, both genes display a similar pattern of expression, showing a sharp increase from 16-cell to 64-cell stage (Fig. 3E). To further explore potential activity of cWnt in 2d, we compared expression patterns of 2d vs 16-cell for components of the cWnt pathway (Fig. 3F). We observed a trend towards up-regulation in 2d of additional pathway components including *Ct-tcf* and *Ct-axin* (Fig. 3F); the latter was proposed to be a cWnt target in *P. dumerilii* (Demilly et al., 2013). This suggests that cWnt signaling is active in 2d as many cWnt components are expressed and a target gene is up-regulated in this blastomere. Additionally, we found the ligand *Ct-nodal* to be enriched in whole-embryos. To understand how this might relate to the role of Nodal/Activin in organizing activity from 2d in *C. teleta*, we used normalized counts data from the 2d vs. 16-cell dataset and identified components of the TGF-β superfamily of pathways (Fig. S4). We observed that most are enrichede in 2d, including a trend for enrichment of the inhibitors *Ct-follistatin, Ct-smad6/7* and the ligand *Ct-bmp3* (Fig. S4). In line with previous evidence on BMP and Nodal/Activin using functional data in *C. teleta* (Lanza & Seaver, 2018, 2020b; Webster et al., 2021; Webster & Meyer, 2024), we propose that 2d may produce and respond to BMP signals based on previous reports of expression of pSMAD1/5/8 (Webster et al., 2021) and produce Nodal/Activin signals based on Nodal expression in this blastomere (Lanza & Seaver, 2020b). This confirms the involvement of the somatoblast 2d as the embryonic organizer and the central role of TGF-β signaling in this process as observed broadly in metazoans.

### Conditional signaling involved in specification of VNC fate

Unlike the brain, which forms in the presence of other blastomeres, the ability of 2d to form the VNC can be repressed or rescued depending on the surrounding blastomeres at the 16-cell stage (Carrillo-Baltodano & Meyer, 2017; Webster et al., 2026). Isolation of 2d with other animal micromeres, 1q^1^, 1q^2^ and 2a–c (−2Q) blocks VNC formation but not in the presence of any one vegetal second-quartet macromere (2A, 2B, 2C or 2D) (Fig. 4A) (Carrillo-Baltodano & Meyer, 2017; Webster et al., 2026). This suggests the existence of pro-neural signals in the vegetal macromeres and anti-neural signals in micromeres at the 16-cell stage. To identify potential pro-neural signals in the 2ACD macromeres, we compared the transcriptomic profiles of 16-cell and 16-cell -2ACD embryos. We found a limited number of differentially-enriched genes in whole-embryos, including *Ct-gataB1,* which is known to be expressed in the 2Q macromeres (Boyle & Seaver, 2008) and *Ct-taleVIIIC*. In contrast, dozens of transcripts were enriched in -2ACD embryos, including *TALE* (*Ct-taleXVIII, Ct-taleIVA, Ct-taleIV-like*) and *FOX* (*Ct-foxq2B, Ct-foxL2/3, foxP1, foxL1*) genes (Fig. 4B, Table S2). From this list, we found *Ct-APC* to be enriched in partial embryos (Fig. 4B), which is a negative regulator of cWnt signaling in other animals. Furthermore, we found gene sets related to cell fate commitment and determination in enriched transcripts in -2ACD embryos (Fig. 4C). To identify potential pro-neural signals and master regulators, we performed a motif enrichment analysis similar to the comparisons above. In enriched transcripts in -2ACD partial embryos, we found four motifs including one with a binding site for FOX (68/80, e-value = 2.5E-002) (Fig. S5). In 16-cell embryos, we found motifs with binding sites for different transcription factors containing an IRF (32/32, e-value = 5.9E-003) or a bZIP-MAF domain (29/32, e-value = 2.6E-003) (Fig. S6). Transcription factors that are associated with those domains are highly expressed during spiral cleavage (Fig. 4D). All in all, our data suggest asymmetrical segregation of TALE and FOX transcription factors along the animal-vegetal axis at the 16-cell stage suggesting a role in cell fate specification.

**Figure 4.**
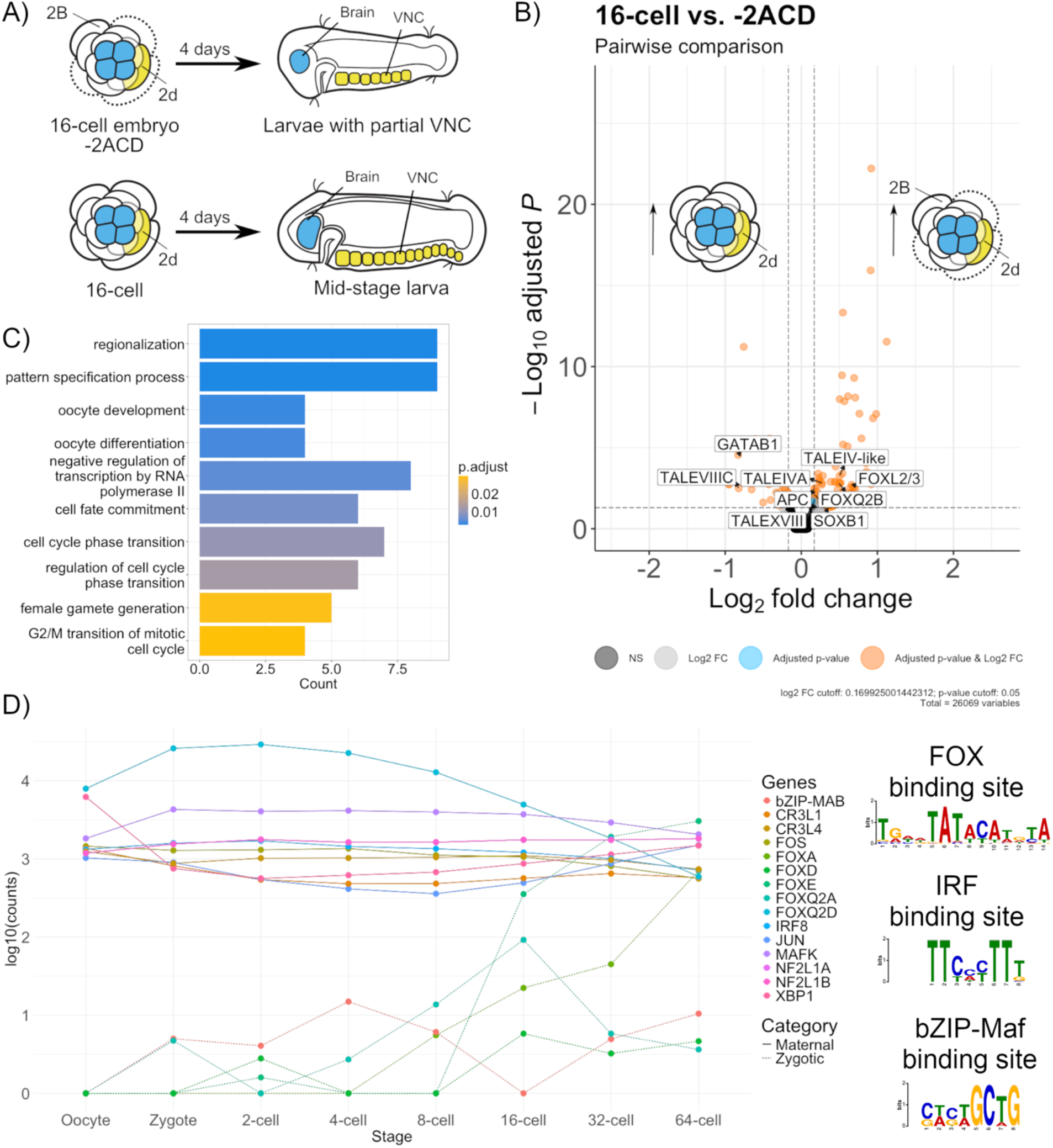
TALE and FOX transcription factors are asymmetrically enriched at 16-cell stage. A) Experimental design for transcriptomic analysis. Top: Removal of 2A, 2B and 2C macromeres (dotted lines) to isolate the animal micromeres (1q daughters + 2q) and macromere 2B (−2ACD isolates). This partial embryo leads to the formation of a reduced VNC. Bottom: 16-cell embryo, with the somatoblast 2d labeled in yellow (left), forming the VNC in a stage 6 larva after ∼4 days (right). B) Volcano-plot showing down-(left) and up-regulated (right) genes in 2q+2B, including differential expression of *FOX* and *TALE* genes. Thresholds for significant differential expression and legends are as in Fig. 2B. C) Bar plot showing the top 10 significantly enriched gene ontology terms (adjusted p-value < 0.05) in genes up-regulated in 16-cell -2ACD embryos. Legend as in Fig. 3D. D) Left: Transcriptomic expression during early development of candidates with a FOX, IRF, and bZIP-Maf DNA binding domain. Right: Consensus binding sites for aforementioned domains found in motifs enriched in -2ACD vs 16-cell.

We previously found that VNC fate is sensitive to repression during the time window between the 16-cell to 32-cell stage (∼2 hours at 21 °C) as we only observed full repression of the VNC at 16-cell (−2Q embryos) but not at 32-cell stage (−3Q) (Fig. 5A) (Carrillo-Baltodano & Meyer, 2017; Webster et al., 2026). Based on this, we performed the following isolations -2Q + 2h (no VNC), -3Q (partial VNC), and whole 32-cell stage embryos (full VNC) from the same brood and performed pairwise comparisons, with the goal of identifying transcriptomic changes occurring at 16-cell to 32-cell stage.

**Figure 5.**
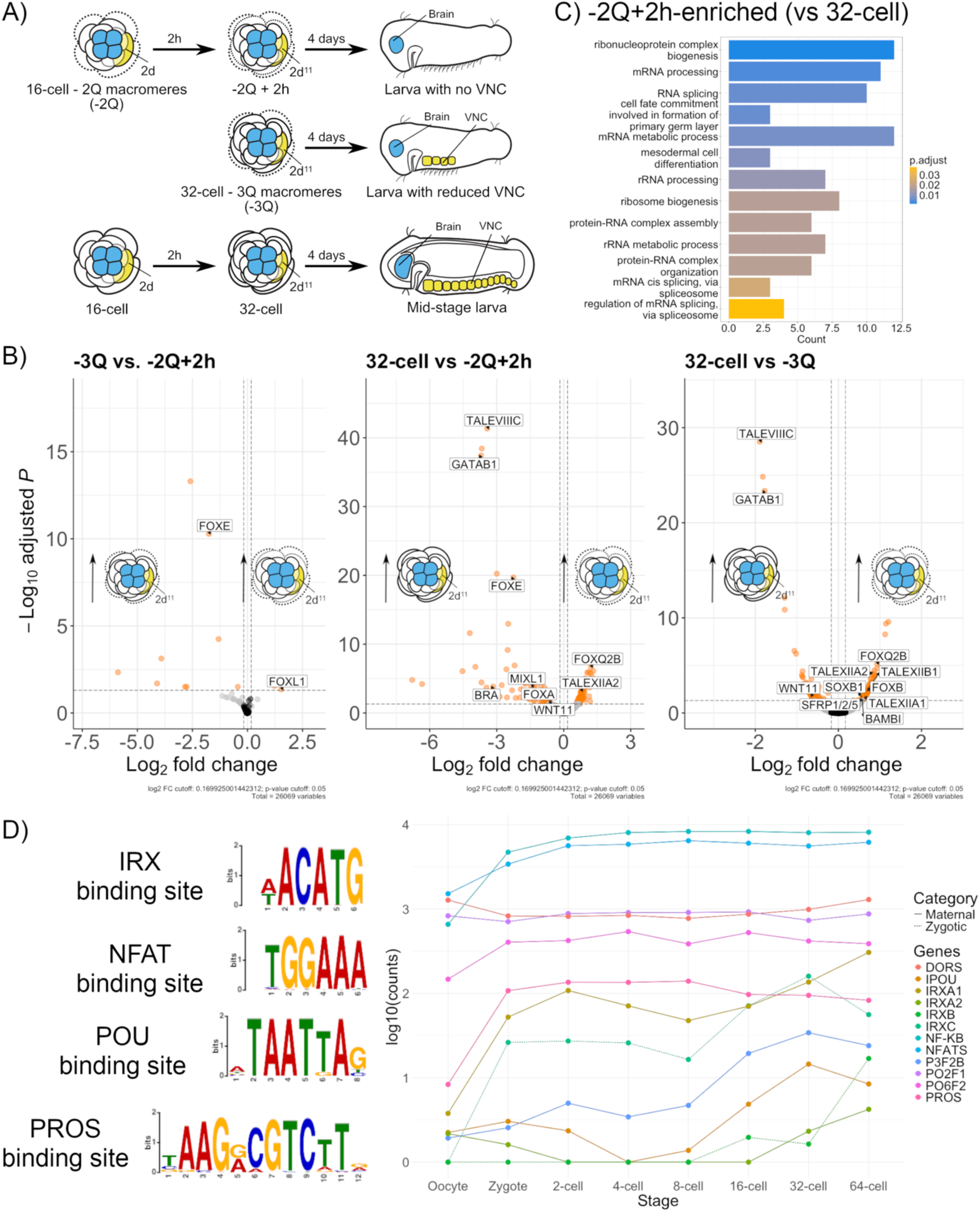
TALE and FOX transcription factors and canonical Wnt signaling pathway components are asymmetrically enriched at 32-cell stage. A) Experimental design for transcriptomic analysis. Top: Removal of 3Q macromeres at 32-cell stage (left) results into a larva having a reduced VNC (right). Middle: Removal of 2Q macromeres at 16-cell stage (left) develops into a partial embryo at 32-cell stage (middle), which then results into a larva missing a VNC (right). Bottom: 16-cell embryo with the somatoblast 2d labeled in yellow (left), 32-cell embryo with the blastomere 2d^1^ in orange (middle), that then develops into a stage 6 larva with a complete ventral nerve cord (VNC, right). B) Volcano-plot showing down-(left) and up-regulated (right) genes in three different pairwise comparisons shown: -2Q+2h vs 3q, -2Q+2h vs 32-cell, -3Q vs 32-cell. Differentially-enriched genes include *TALE, FOX*, and cWnt components. Thresholds for significant differential expression and legends are as in Fig. 2B. C) Bar plot showing significantly enriched gene ontology terms (adjusted p-value < 0.05) in genes up-regulated in 16-cell -2ACD embryos. Legend as in Fig. 3D. D) Left: Consensus binding sites for IRX, NFAT, POU and PROS found in motifs enriched in transcripts from all pairwise combinations as in Fig. 5A. Right: Transcriptomic expression during early development of candidates with aforementioned domains.

Based on previous results (Fig. 1C), we sought to identify targets of anti-neural signaling affecting VNC fate by comparing the transcriptomic profiles of isolates where macromeres were removed within a 2h time frame (−2Q + 2h vs -3Q). In this comparison, we observed that *Ct-foxL1* and *Ct-bsx* were enriched in -2Q + 2h embryos (Fig. 5B), while *Ct-foxE* and *Ct-lhx1/5* were enriched in -3Q (Fig. 5B). Subtle differences between treatments in timing and number of blastomeres show that timing during spiral cleavage is crucial for signaling events and the expression of blastomere-specific transcription factors.

To identify signaling events and transcription factors involved in neural fate specification in the ventral nerve cord, we compared the transcriptomic profiles of -2Q + 2h vs. 32-cell, and - 3Q vs. 32-cell. As observed for all pairwise comparisons at 8-cell and 16-cell stage, differential expression of *FOX* and *TALE* genes remained consistent in these comparisons as well (Fig. 5B, Table S5). In both -2Q+2h vs 32-cell and -3Q vs 32-cell, we observed a similar set of *FOX* and *TALE* genes as up-(*Ct-foxq2B, Ct-foxB2, Ct-foxL1, Ct-taleXIIA1, Ct-taleXIIA2, Ct-taleXIIB1)* and down-regulated (*Ct-taleVIIIC)* (Fig. 5B, Table S2). Furthermore, we found a gene set enriched involved in mesodermal cell differentiation (Fig. 5C). In addition, we also identified transcripts that were unique to one comparison. We found a smaller set of enriched transcripts when comparing -3Q vs 32-cell (116 in partial embryos and 86 in whole embryos) and -2Q+2h vs 32-cell (158 in partial embryos and 132 in whole embryos) (Fig. 5C). These additional transcripts included other FOX and TALE genes. Across both comparisons, we found *Ct-sfrp1/2/5B* and *Ct-soxB1* to be consistently animal, with the latter known to be involved in neurogenesis later in development in *C. teleta* (Sur et al., 2017, 2020). In contrast, aside from *Ct-taleVIIIC,* we found that *Ct-wnt11* and *Ct-gataB1* were consistently vegetal (Fig. 5B), with the latter known to be expressed in 2Q and 3Q macromeres in *C. teleta* (Boyle & Seaver, 2008). Differential expression of *Ct-sfrp1/2/5B* and *Ct-wnt11* along the animal-vegetal axis suggests a potential Wnt signaling gradient similar to that observed across metazoans (Marlow et al., 2014). Altogether, we suggest that a combination of *FOX* and *TALE* genes, Wnt signaling and additional transcription factors may be important for neural fate specification and blastomere identity in *C. teleta*.

To identify potential pro- and anti-neural signals and master regulators underlying VNC specification, we performed a motif enrichment analysis in sets of DEGs in all comparisons to identify master regulators within the same species that might control gene expression across the animal-vegetal axis. When comparing -2Q+2h vs 32-cell, we found four motifs in transcripts enriched in -2Q+2h embryos, including one with a binding site for IRX (134/158, e-value = 1.6E-007), and other classes of transcription factors (Fig. 5D). All possible upstream transcription factors containing DNA-binding domains from the previoys analysis are expressed at 32-cell stage at different levels (Fig. 5D). However, their functional role remains to be tested.

Altogether, pairwise comparisons from all isolations at 8-, 16-, and 32-cell stage suggest a fine-tuned control of cell-cell signaling and transcription factor dynamics led by *TALE* and *FOX* genes underlying neural fate specification and blastomere identity.

## Discussion

Using blastomere isolations and RNA-sequencing, we have provided the first reported model of neural specification in Spiralia and the molecular profiles of partial embryos at different stages during spiral cleavage in an annelid. We have also provided the first reported transcriptomic profile of the organizer 2d in Spiralia, which can be used for cross-species comparison with organizers in other annelids and non-annelid spiralians. While functional data is needed to confirm the role of *FOX* and *TALE* transcription factors and cWnt signaling and sample sizes were limiting due to material availability, these data can still provide major insights into a putative molecular mechanisms underlying neural specification and blastomere identity in Spiralia.

### Brain specification

In multiple spiralian species, the 1q micromeres form the ectoderm including neural tissue of the episphere and the prototroch ciliary band. This is the case for the annelids *Capitella teleta* (Fig. 1A-B) and *Platynereis dumerilii,* the platyhelminth *Hoploplana inquilina,* the nemertean *Cerebratulus lacteus* and the mollusks *Crepidula fornicata* and *Patella vulgata* (Ackermann et al., 2005; Boyer et al., 1998; Dictus & Damen, 1997; Hejnol et al., 2007; Henry & Martindale, 1998; Meyer & Seaver, 2010). This indicates that 1q micromeres have similar contributions to the body plan across multiple phyla. In both annelids *Owenia fusiformis* and *C. teleta*, a very limited number of genes are differentially-expressed between 4-cell vs 8-cell embryos (Liang et al., 2025). This is consistent with the limited number of differentially-enriched genes we found in 1q vs 8-cell, suggesting that only a few transcription factors are asymmetrically segregated across the animal-vegetal axis before the organizer and somatoblast 2d is born. Expression data during spiral cleavage in *O. fusiformis* (Seudre, Martín-Zamora, et al., 2022) and the mollusk *Nipponacmea fuscoviridis* (Morino & Yoshikawa, 2023) are similar to our findings in *C. teleta* where *Ct-foxq2A* and *Ct-foxq2B* are enriched in 1q micromeres, suggesting that *FOXQ2* paralogs are localized to the animal pole in *C. teleta*. Similar patterns in annelids and mollusks suggest that the asymmetric segregation of *FOXQ2* mRNA towards the 1q micromeres may be ancestral for Spiralia. This also suggests that these genes potentially confer blastomere identity to 1q as they are enriched in this quartet of blastomeres. Altogether, we hypothesize that this could be a conserved mode of cell fate specification in Spiralia.

Beyond spiralians, *FOXQ2* has been widely studied in other taxa, showing repeated anteriorly-localized expression across metazoans, and has been hypothesized to have an ancestral role in specification of anterior neural fate in bilaterians (Gattoni et al., 2025; Marlow et al., 2014; Morino & Yoshikawa, 2023; Santagata et al., 2012). In the cnidarian *Clytia haemispherica,* overactivation of Wnt/β-catenin signaling leads to the repression of aboral markers including *foxQ2a* (Chevalier et al., 2006; Momose et al., 2008). In the echinoderm *Strongylocentrotus purpuratus*, Wnt/β-catenin signaling from the vegetal pole restricts the expression of *Sp-foxQ2* and specification of the neurogenic ectoderm to the animal pole (Cui et al., 2014; Range & Wei, 2016; Yaguchi et al., 2008). In the insect *Tribolium castaneum,* Wnt/β-catenin represses *Tc-six3* and *Tc-foxQ2,* which are upstream in the anterior GRN (Kitzmann et al., 2017). All in all, across metazoans, maternal/zygotic *foxQ2* expression is spatially restricted by Wnt/β-catenin signaling, which might explain its conservation in *C. teleta*. With our results and in context with data from other taxa, we propose that *C. teleta* has a similar ancestral mode of specification for the anterior neuroectoderm, with *Ct-foxq2A* and *Ct-foxq2B* acting as possible brain neural determinants. All in all, this suggests that the brain (or anterior neural tissue) is homologous across Spiralia.

### Specification of the somatoblast 2d

Differences in modes of early development and life history lead to major distinctions in body plan formation. In multiple annelids, the VNC arises from 2d, including *Amphitrite ornata, Nereis, Scoloplos armiger, Helobdella, Platynereis dumerilii* and *C. teleta* (Fig. 1A, B) (Ackermann et al., 2005; Mead, 1897; Meyer et al., 2010; Meyer & Seaver, 2010; Weisblat et al., 1984). However, outside of Annelida, fate-mapping data in other spiralians do not show a conservation in the formation of trunk CNS tissue from descendants of 2d (Meyer et al., 2010). We observed a unique set of transcription factors enriched in 2d in *C. teleta*, coinciding with the onset of zygotic transcription (Liang et al., 2025). Notably, *FOX* genes were enriched in 2d, including two other paralogs of *FOX*Q2 *(Ct-foxq2C and Ct-fox2D).* In contrast with 1q, we observed enrichment of *TALE* genes in 2d, including *Ct-taleVIIIB1* and *Ct-taleIV-like.* Overall, this supports the involvement of *FOX* and *TALE* genes in cell fate segregation at 16-cell stage (Fig. 5C). Based on its high expression in oocytes compared to other 2d-enriched transcripts, we propose *Ct-foxq2D* as a determinant of 2d (Fig. S3), pointing towards a mechanism similar to 1q *(Ct-foxq2A* and *Ct-foxq2B)*, and potentially distinctive from how the VNC is specified in non-spiralian taxa (Ashe et al., 2000; Delaune et al., 2005). Functional testing of *FOXQ2* paralogs in *C. teleta* would provide additional evidence towards a different mode of neural specification of the VNC in Spiralia, contributing to our understanding of nerve cord evolution in bilaterians (Martín-Durán et al., 2018). In addition, Spiralia-specific TALE homeobox genes have been proposed as important for blastomere specification, with reported blastomere-specific expression in the annelid *Spirobranchus kraussii* (Morino et al., 2017). In *C. teleta,* we found *TALE-VIIIB1* and *TALE-IV-like* enriched in 2d, indicating that SPILE genes could also confer molecular identity to blastomeres in unequal-cleavers. Altogether, these data provide insights into molecular mechanisms underlying blastomere identity, and how *TALE* and *FOX* genes might have a central role during early development in Spiralia.

### Organizing activity in Capitella teleta and Spiralia

Organizing activity in Spiralia is tightly linked to spiral cleavage as organizer cells are specified during early development (Lambert & Nagy, 2001; Seudre, Carrillo-Baltodano, et al., 2022). Across spiralians, the identity of the D-V organizer seems to be restricted mostly to the D-quadrant, as it is shown to be in 2d in *C. teleta* (Amiel et al., 2013), 4d (64-cell stage) in *O. fusiformis* (Seudre, Carrillo-Baltodano, et al., 2022) and the slipper-snail *Crepidula fornicata* (Henry & Perry, 2008), 2d^11^ and 4d in the oligochaete *Tubifex tubifex* (Nakamoto et al., 2011) and 3D (32-cell stage) in the mud snail *Ilyanassa obsoleta* (Lambert & Nagy, 2001). In all species mentioned above, except for *T. tubifex* and *C. teleta*, it has been shown that phosphorylation of ERK1/2 is required to specify the organizer (Henry & Perry, 2008; Lambert & Nagy, 2003; Seudre, Carrillo-Baltodano, et al., 2022). However, in *C. teleta*, phosphorylated ERK1/2 was detected during gastrulation but not cleavage stages despite 2d being identified as the organizer (Amiel et al., 2013). This indicates that a different signaling event may specify the organizer in this annelid, possibly via Wnt signaling based on our 2d-enriched transcripts.

Equal spiral cleavage is considered ancestral for Spiralia (Martín-Durán & Marlétaz, 2020), with BMP having a central role in dorsal-ventral patterning in equal-cleaver annelids (Carrillo-Baltodano et al., 2026). However, in unequal-cleavers, such as *Platynereis dumerilii* and *C. teleta*, BMP has a different role during early development (Carrillo-Baltodano et al., 2026; Webster et al., 2021; Webster & Meyer, 2024). In *C. teleta,* functional experiments showed that Nodal/Activin is the primary D-V organizing signal (Lanza & Seaver, 2020b). Our data here support that finding by showing enrichment of Activin/Nodal ligands in 2d. The identity of the cells responding to Nodal/Activin and whether other pathways act in parallel to establish axes and instruct surrounding blastomeres require further functional work. However, while the role of BMP and Nodal/Activin in D-V axis specification seems to be ancestral in annelids and possibly Spiralia, coupling this process to neural fate specification remains contested as emerging data suggest evolution of a spiralian-specific mechanism controlling neural specification.

### Model for combined conditional and autonomous neural specification in C. teleta

We previously established time windows for specification of the brain and nerve cord in *Capitella teleta* (Fig. 6B). In our current dataset, we found cWnt pathway components enriched along the animal-vegetal axis at the 16-cell and 32-cell stage (Fig. 3-5). For instance, the animal blastomeres (in -2Q + 2h and -3Q) have an enrichment of *Ct-sfrp1/2/5,* while 32-cell embryos display enrichment of *Ct-wnt11.* From all pairwise comparisons, we created a model of how cWnt might operate during neural specification in *C. teleta* (Fig. 6C). First, we propose that *FOXQ2* genes act as maternal determinants for the brain and ventral nerve cord, whose levels can be controlled based on Wnt/β-catenin activity. Then, incorporating data about animal-vegetal differences in β-catenin levels in the annelid *P. dumerilii* (Schneider & Bowerman, 2007), we propose the following model: (1) A gradient of cWnt signaling is established along the animal-vegetal axis of the embryo, with 1q micromeres (anterior neuroectoderm) having lower levels of β-catenin and 2d (trunk neuroectoderm) having higher levels. (2) In the anterior neuroectoderm, cWnt activity remains lower due to the expression of the antagonist *sFRP1/2/5* in the animal pole. (3) In the trunk neuroectoderm, cWnt activity remains higher due to the expression of Wnt11 in the vegetal pole. (4) In the absence of the vegetal macromeres, cWnt signaling is repressed via *sFRP1/2/5* from the animal micromeres, leading to lower levels of β-catenin and repression of neural fate specification in the trunk.Meanwhile, β-catenin levels remain unchanged in the animal pole, resulting in unaffected head neuroectoderm. Altogether, this model where *FOXQ2* are neural determinants and Wnt signaling promotes neural fate via ligand/antagonist expression also suggests the conservation of a FoxQ2-Wnt GRN that is also involved in the trunk neuroectoderm in *C. teleta* during spiral cleavage (Fig. 6D). This GRN is involved in specification of anterior neuroectoderm in multiple metazoans (Marlow et al., 2014), suggesting redeployment of an anterior GRN during trunk neuroectoderm specification in Spiralia.

**Figure 6.**
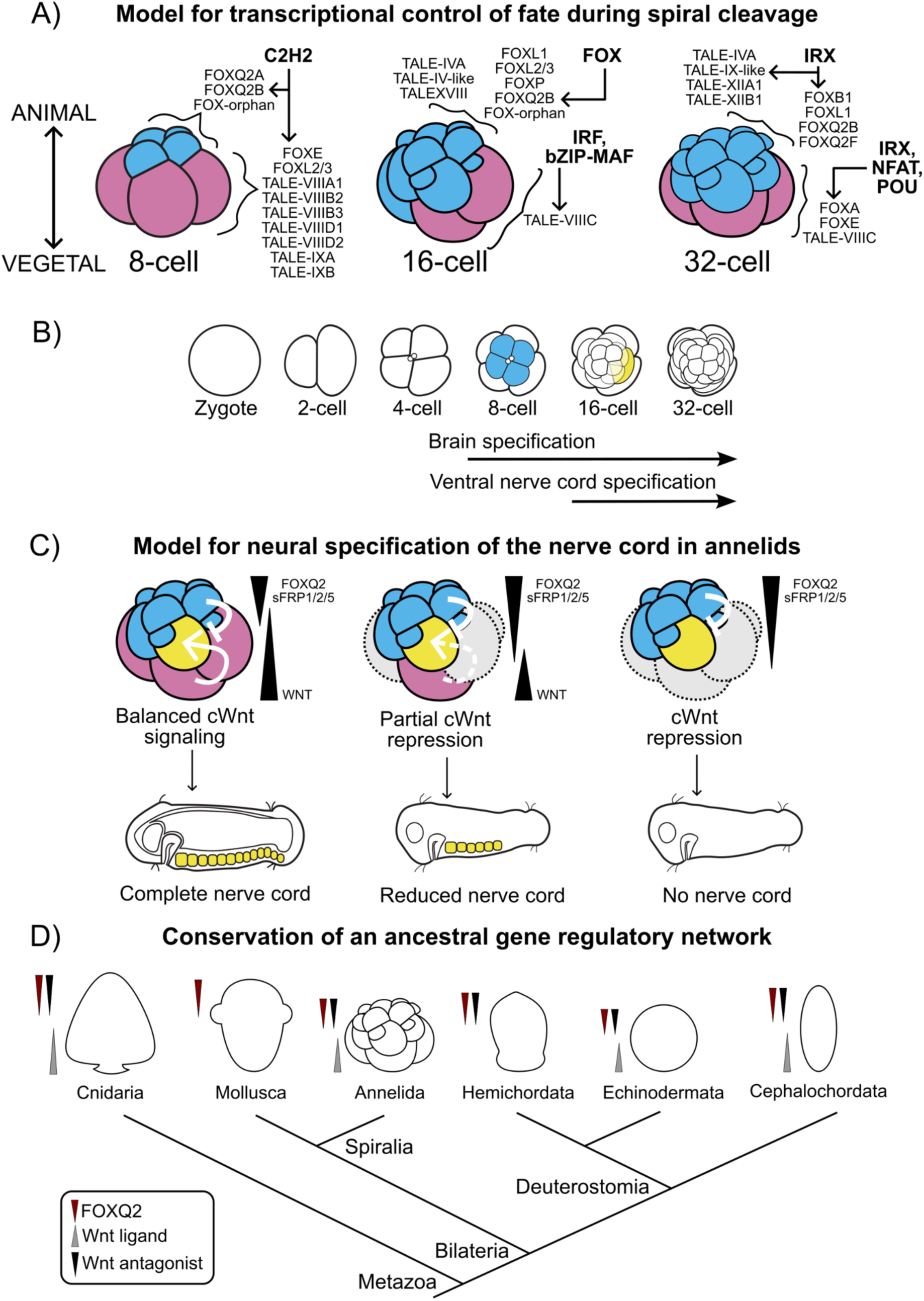
Model for cell fate specification during spiral cleavage. A) Model for transcriptional control of fate during spiral cleavage based on the asymmetrical enrichment of mRNAs along the animal-vegetal (A-V) axis and proposed master regulators based on RNA-seq and binding site analysis. Across all stages, consistent enrichment of similar *FOX* and *TALE* genes are observed across the embryonic axis, indicating a fine-tuned transcriptional control of cell fate segregation during spiral cleavage of *C. teleta*. B) Time window for cell fate specification and brain and ventral nerve cord (VNC) specification in *Capitella teleta.* Brain specification occurs at 8- to 16-cell stage, where blastomeres that originate the brain are born. VNC specification occurs at 16- to 32-cell stage, where the blastomere that originates the VNC is born and when pro- and anti-neural signals occur based on previous blastomere isolation experiments. C) Our model for neural fate specification of the VNC in annelids. In a whole-embryo, a balance of activation/repression of canonical Wnt signaling (cWnt) leads to the formation of a complete VNC (left). In a partial embryo where the macromere 2B is present, a partial repression of cWnt is observed which leads to a reduced nerve cord (middle). In a partial embryo where all macromeres are removed, a higher repression of cWnt leads to the repression of nerve cord formation (right). D) Phylogeny indicating the conservation of an ancestral gene regulatory network indicating gradients of expression of FoxQ2-Wnt in multiple taxa, also observed now in *C. teleta* during early embryogenesis. Adapted from Marlow et al., (2014).

### Establishment of blastomere identity during spiral cleavage

Spiral cleavage is a conserved developmental program present in at least eight phyla including annelids and mollusks (Martín-Durán & Marlétaz, 2020). Data from annelids and mollusks suggest that ERK1/2, BMP and Nodal/Activin are active during spiral cleavage and are central for the establishment of body axes in the future larva (Carrillo-Baltodano et al., 2026; Henry & Perry, 2008; Lambert & Nagy, 2001; Lanza & Seaver, 2020b; Seudre, Carrillo-Baltodano, et al., 2022). However, data related to how blastomere identities are conferred during spiral cleavage before organizing activity occurs has been explored only in mollusks (Morino et al., 2017; Morino & Yoshikawa, 2023; Phuangphong et al., 2021). In the equal spiral-cleaving limpet *Nipponacmea fuscovidiris*, five lineage-specific *TALE* homeobox genes known as *SPILE* genes were functionally tested and found to comprise a gene regulatory network (GRN) during early cleavage(Morino et al., 2017; Morino & Yoshikawa, 2023). In the unequal spiral-cleaving *Mytilosepta virgata*, an expansion of *TALE* genes was observed, and all were found to be expressed in a quadrant- or blastomere-specific manner, suggesting a role in blastomere identity (Phuangphong et al., 2021). In our data, we found differential enrichment across the animal-vegetal axis of different *SPILE* genes in *C. teleta* at 8-cell, 16-cell and 32-cell stages, suggesting the existence of a similar GRN in the annelid with the largest expansion of SPILE genes reported (Barton-Owen et al., 2018). Across all stages examined, we found enrichment of a stage-specific set of SPILE genes along the animal-vegetal axis, with a few overlapping genes at consecutive stages (Fig. 6A). Based on this, we hypothesize that both *TALE* and *FOX* genes are involved in blastomere identity in *C. teleta* (Fig. 6D), but confirmation of this requires gene expression and functional studies. In connection with the role of BMP signaling in establishing blastomere identity in *C. teleta* and *P. dumerilii* (Carrillo-Baltodano et al., 2026; Webster et al., 2021), we hypothesize that BMP signaling regulates expression of *FOX* and *TALE* genes in the animal pole to confer blastomere identity.

In conclusion, by combining blastomere isolations and RNA-sequencing, we showed an asymmetrical segregation of mRNA across the animal-vegetal axis of the embryo during spiral cleavage in a temporal manner, identified markers for the blastomeres originating the CNS, and proposed a model for neural specification in *C. teleta* (Fig. 6C). We suggest that a similar gene hierarchy involving both lineage-specific and conserved transcription factors is necessary to assign blastomere identity during spiral cleavage, and an ancestral FoxQ2-Wnt GRN and balance of cWnt signaling is central for neural specification instead of inhibition of BMP signaling in Annelida.

## Materials and Methods

### Animal care

A laboratory colony of *Capitella teleta* adults and earlier stages is maintained all-year in glass finger bowls with 32–34 ppt artificial seawater (ASW) and mud collected from Rye, NH 2–3 times a year (Blake et al., 2009). Adult bowls are kept at 19 °C as described elsewhere (Seaver et al., 2005). For these experiments, embryos were collected and raised in 0.22 µm-filtered ASW with 60 µg/mL penicillin and 50 µg/mL streptomycin (P/S FASW) (Sigma Aldrich). Early-cleavage stages were obtained by dissecting the broods with Dumont #5 forceps and identified visually based on its stereotypical cleavage program as described previously (Meyer et al., 2010). All embryos were collected in gelatin-coated dishes at 15 or 19 °C until treatment.

### Blastomere isolation

Embryos at the selected early-cleavage stage were incubated in Ca^2+^ Mg^2+^ -free ASW for 5 min in 24-well gelatin-coated dishes, followed by an incubation for 5 min in 1% Protease (Sigma-Aldrich cat# P6911) and 1% Sodium thioglycolate (Sigma-Aldrich cat# T0632) as described elsewhere (Carrillo-Baltodano & Meyer, 2017). Embryos were washed three times with P/S FASW post-treatment. Blastomeres were removed as desired using an eyelash attached to a glass pipette. For 2d isolations, this procedure was done as above to isolate 1D, and then blastomeres were maintained in P/S FASW until they divided (∼40 min, 2D and 2d), and 2D was removed (Webster et al., 2026). All isolations performed for pairwise comparisons are specified in Table S5. Untreated controls and isolates were transferred to 35 mm gelatin-coated dishes with P/S FASW until they were ready to be lysed for RNA extraction.

### RNA extraction

The Arcturus Picopure™ RNA Isolation Kit (Thermo Fisher Scientific, cat# KIT0204) was used to isolate RNA as it is designed for low-input samples. Post-isolations, embryos and cells in minimum P/S FASW were lysed, incubated at 42 °C and stored at -80 °C until further processing as per manufacturer’s instructions. Samples were processed as per manufacturer’s instructions. DNase treatment was performed using the RNase-free DNase Set (Qiagen, cat# 79254). RNA was quantified using the Qubit™ RNA High Sensitivity kit (Thermo Fisher Scientific, cat# Q32852) and sent for library preparation and sequencing to Novogene Corporation (Sacramento, CA). Samples with expected low yield (<100 ng/µL) were processed using the ultra low-input pipeline.

### Library preparation and quality control

A low-input RNA-sequencing approach was requested to Novogene Corporation. A minimum of 100 ng was required to be sent per sample to perform mRNA enrichment, RNA fragmentation, reverse transcription, library preparation and paired-end 150 bp (PE150) sequencing. For those samples going to the ultra low-input pipeline (ULI), the minimum required was 10 ng total. Samples were assessed using Nanodrop (concentration in ng/µL) and Agilent Bioanalyzer (RNA Integrity Number). ULI-processed samples were processed for library preparation using SMART-Seq® v4 Ultra® Low Input RNA Kit and proceeded to PE150 sequencing. Major differences between low-input and ultra low-input were based on the experimental design (time constraints based on brood size) and yield per isolate (Table S5). Sequencing for all samples was done in Illumina NovaSeq 6000. Quality of the output was assessed based on clean and total reads and base error rate (Table S5, S5).

### Bioinformatic processing of raw sequencing data

Raw-data quality was assessed using FastQC 0.12.0 (Andrews, 2023). Reads with low-quality bases were removed using Trimmomatic v.036 (Bolger et al., 2014). For ULI samples, an additional step to remove Nextera Transposase Sequences was done using TrimGalore (Krueger et al., 2023). Filtered reads were mapped to a chromosome-level assembly of the genome of *Capitella teleta* (Davies et al., 2026) using STAR v2.7.11b (Dobin et al., 2013) showing >69% mapping rate for all samples (Table S5). Counts were generated using featureCounts (Liao et al., 2014). Analysis of differential gene expression was done separately for each set of proposed comparisons using the R package DEseq2 v1.42.1 (Love et al., 2014). An adjusted p-value of less than 0.05 and a Log2fold value of greater or lower than 0.17 was used to determine significance in differential gene expression. A lower threshold for differential expression was applied to detect subtle but significant changes in gene expression across samples. Principal component analysis was performed in R v4.4.0 and plotted using ggplot2. Hierarchical clustering was also performed in R v4.4.0 and plotted using pheatmap. Expression data was visualized using a volcano plot, done using the R package EnhancedVolcano available in Bioconductor. Pairwise comparisons were done to assess differential expression across different conditions during early-cleavage (Table S5).

### Identification of transcription factors

Using an existing annotation (Davies et al., 2026), we identified all transcription factors present in the list of differentially-enriched genes (DEGs). Identity was confirmed using the Conserved Domain Database and BLAST+ (Camacho et al., 2009; Marchler-Bauer et al., 2003). *TALE* and *FOX* genes were filtered from significant differentially-enriched genes and reannotated using a modified version of a pipeline for homeobox gene annotation (Zwarycz et al., 2016).

### Gene ontology analysis

Signaling pathway components were identified using the pre-existing annotation for this species (Davies et al., 2026). A package for *Capitella teleta* was generated based on GO annotations using the R package AnnotationForge (Carlson & Pagès, 2026). Over-representation of Gene Ontology (GO) terms from differentially-enriched genes per treatment were identified using clusterProfiler and visualized using the package enrichplot within that environment (Yu et al., 2012).

### Transcription factor binding site analysis

Using bedtools, we extracted 2.5kb upstream and 250 bp downstream from the transcription start for each gene enriched per treatment (Quinlan & Hall, 2010). Using these sequences, a motif enrichment search was done initially in FIMO and then in MEME (nmotifs = 10) with a range of 6-30 bp per motif (p-value < 0.05) within the MEMEsuite environment v5.5.9 (Bailey et al., 2015). To identify known binding sites within the motif found, we used TOMTOM with a threshold of e-value < 1 (Gupta et al., 2007). Candidates were then identified based on PFAM DNA-binding domains in the proteome of *C. teleta* and its mRNA expression during spiral cleavage was obtained using the developmental transcriptome existing for this species (Liang et al., 2025).

## Author contributions

Conceptualization: J.D.-S., N.P.M.; Methodology: J.D.-S., N.P.M.; Software: J.D.-S., J. T., N.P.M.; Investigation: J.D.-S., N.P.M.; Writing – original draft: J.D.-S.; Manuscript revision: N. P. M. Visualization: J.D.-S., N.P.M.; Supervision: N.P.M.; Project administration: N.P.M.; Funding acquisition: J.D.-S., J.T., N. P. M.

## Acknowledgements

We would like to thank Dr. José María Martín-Durán at Queen Mary’s University of London for providing early access to the genome assembly and developmental transcriptome of *Capitella teleta* and for providing valuable feedback in motif enrichment and transcription factor footprinting analysis. We would also like to thank Alfredo Cubas Hidalgo, Lizanne Otañez-Vargas and Thomas Roehl for their assistance for shipment of samples for RNA-sequencing, and Dr. Alexander Bradshaw for his continuous help with sample shipment, data analysis and advice during the first steps of manuscript writing.

## Data availability

Coding resources and metadata can be accessed through the GitHub repository page (https://github.com/npmeyer-capitella/Annelid-Blastomere-RNAseq.git). Raw reads will be deposited in SRA/NCBI.

