## Supplementary Figures and Tables for "Unraveling molecular mechanisms for specification of blastomere identity and neural fate in the annelid *Capitella teleta*"

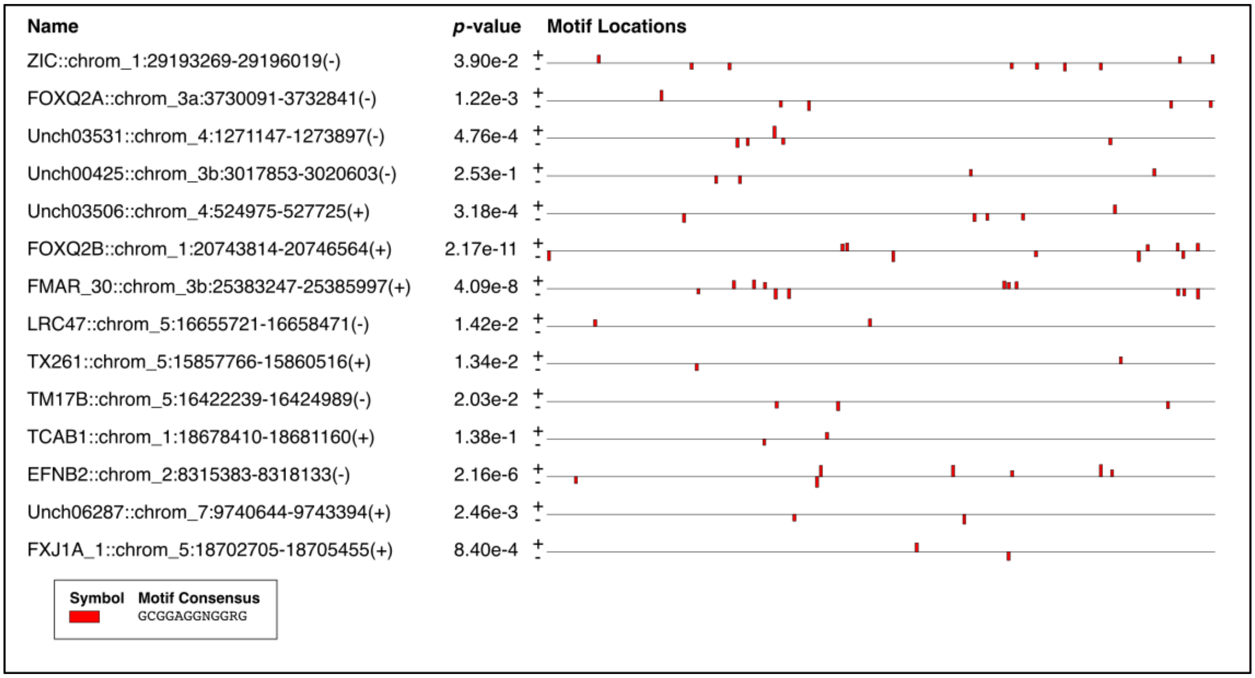


**Figure S1.**

Motif enrichment analysis via MEME for -1Q differentially-enriched transcripts. Column 1 represents the name of the gene, assigned based on their localization in the chromosome after extraction using bedtools. Column 2 is the p-value, representing the probability of a random motif in the sequence to match the motif. Column 3 shows the localization of the motifs. Enriched motif is displayed in red visualized over the 2750 bp region to observe specific patterns of clustering across different genes.

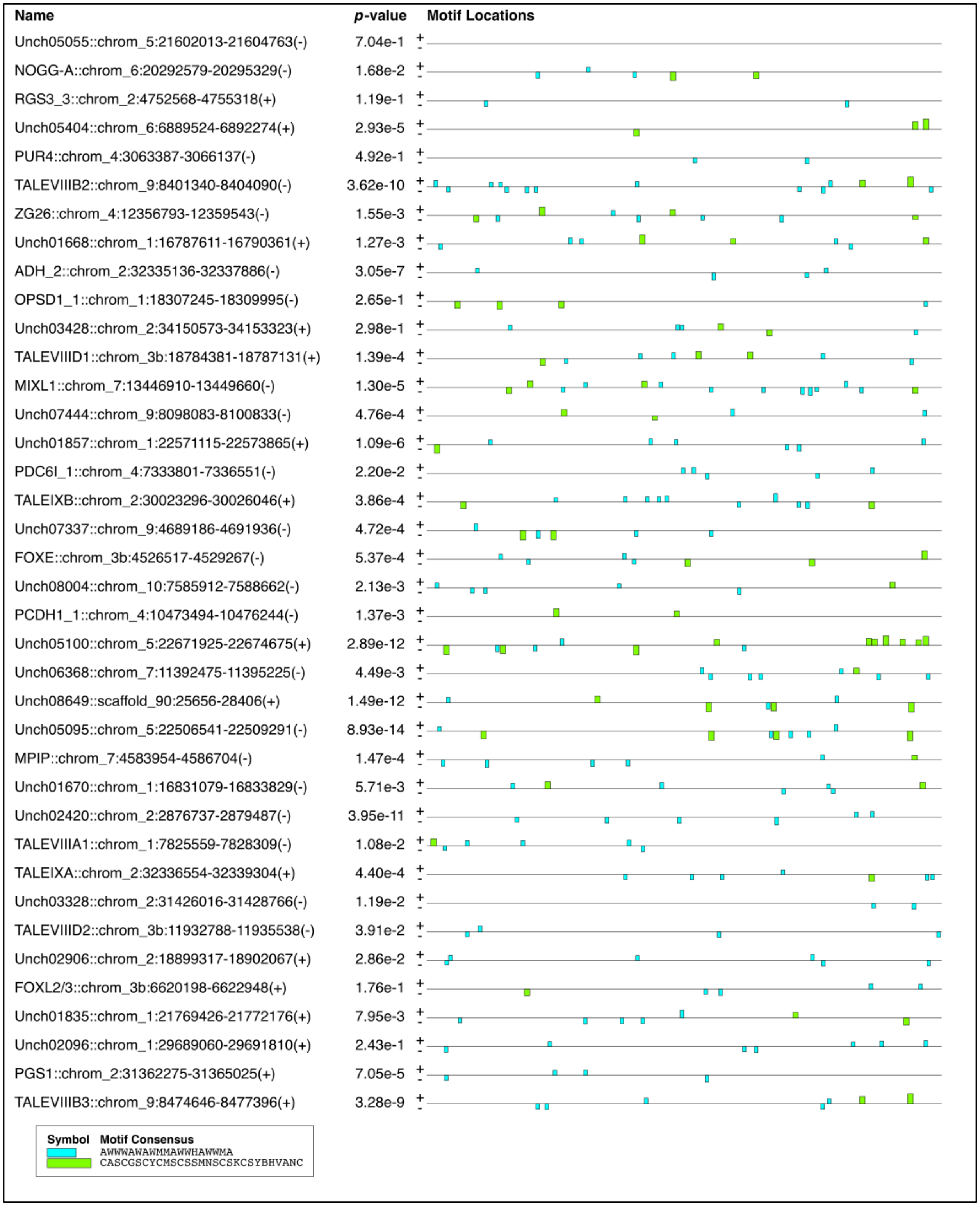
Name
p-value
Motif Locations
Unch05055::chrom_5:21602013-21604763(-) 7.04e-1
NOGG-A::chrom_6:20292579-20295329(-) 1.68e-2
RGS3_3::chrom_2:4752568-4755318(+) 1.19e-1
Unch05404::chrom_6:6889524-6892274(+) 2.93e-5
PUR4::chrom_4:3063387-3066137(-) 4.92e-1
TALEVIIIB2::chrom_9:8401340-8404090(-) 3.62e-10
ZG26::chrom_4:12356793-12359543(-) 1.55e-3
Unch01668::chrom_1:16787611-16790361(+) 1.27e-3
ADH_2::chrom_2:32335136-32337886(-) 3.05e-7
OPSD1_1::chrom_1:18307245-18309995(-) 2.65e-1
Unch03428::chrom_2:34150573-34153323(+) 2.98e-1
TALEVIIID1::chrom_3b:18784381-18787131(+) 1.39e-4
MIXL1::chrom_7:13446910-13449660(-) 1.30e-5
Unch07444::chrom_9:8098083-8100833(-) 4.76e-4
Unch01857::chrom_1:22571115-22573865(+) 1.09e-6
PDC6I_1::chrom_4:7333801-7336551(-) 2.20e-2
TALEIXB::chrom_2:30023296-30026046(+) 3.86e-4
Unch07337::chrom_9:4689186-4691936(-) 4.72e-4
FOXE::chrom_3b:4526517-4529267(-) 5.37e-4
Unch08004::chrom_10:7585912-7588662(-) 2.13e-3
PCDH1_1::chrom_4:10473494-10476244(-) 1.37e-3
Unch05100::chrom_5:22671925-22674675(+) 2.89e-12
Unch06368::chrom_7:11392475-11395225(-) 4.49e-3
Unch08649::scaffold_90:25656-28406(+) 1.49e-12
Unch05095::chrom_5:22506541-22509291(-) 8.93e-14
MPIP::chrom_7:4583954-4586704(-) 1.47e-4
Unch01670::chrom_1:16831079-16833829(-) 5.71e-3
Unch02420::chrom_2:2876737-2879487(-) 3.95e-11
TALEVIIIA1::chrom_1:7825559-7828309(-) 1.08e-2
TALEIXA::chrom_2:32336554-32339304(+) 4.40e-4
Unch03328::chrom_2:31426016-31428766(-) 1.19e-2
TALEVIIID2::chrom_3b:11932788-11935538(-) 3.91e-2
Unch02906::chrom_2:18899317-18902067(+) 2.86e-2
FOXL2/3::chrom_3b:6620198-6622948(+) 1.76e-1
Unch01835::chrom_1:21769426-21772176(+) 7.95e-3
Unch02096::chrom_1:29689060-29691810(+) 2.43e-1
PGS1::chrom_2:31362275-31365025(+) 7.05e-5
TALEVIIIB3::chrom_9:8474646-8477396(+) 3.28e-9
Symbol
Motif Consensus
AWWWAWAWMMAWWHAWWMA
CASCGSCYCMSCSSMNSCSKCSYBHVANC

**Figure S2.**

Motif enrichment analysis via MEME for 8-cell differentially-enriched transcripts. Column 1 represents the name of the gene, assigned based on their localization in the chromosome after extraction using bedtools. Column 2 is the p-value, representing the probability of a random motif in the sequence to match the motif. Column 3 shows the localization of the motifs. Enriched motif is displayed in red visualized over the 2750 bp region to observe specific patterns of clustering across different genes.

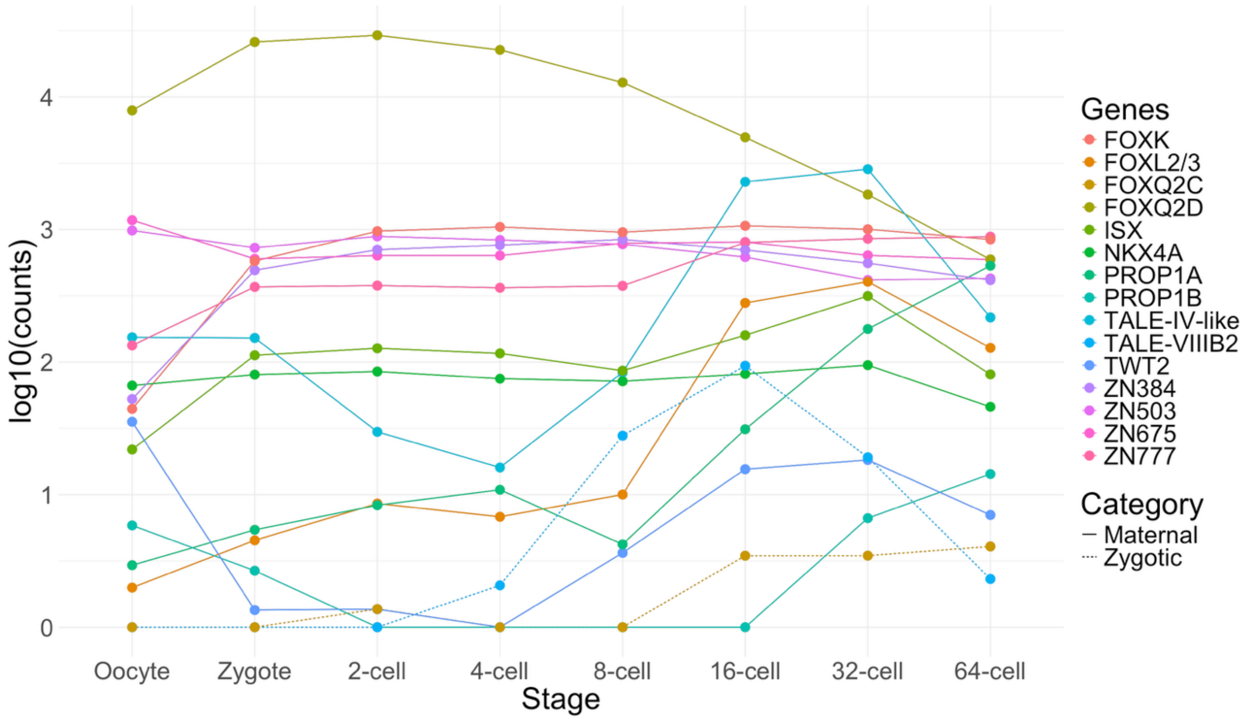


**Figure S3.**

Transcript-level normalized counts during spiral cleavage in *Capitella teleta* for all transcription factors that are enriched in the somatoblast 2d using the developmental transcriptome available for this species (Liang et al., 2025). *Ct-foxq2D* is highly expressed maternally and is proposed as a determinant for 2d.

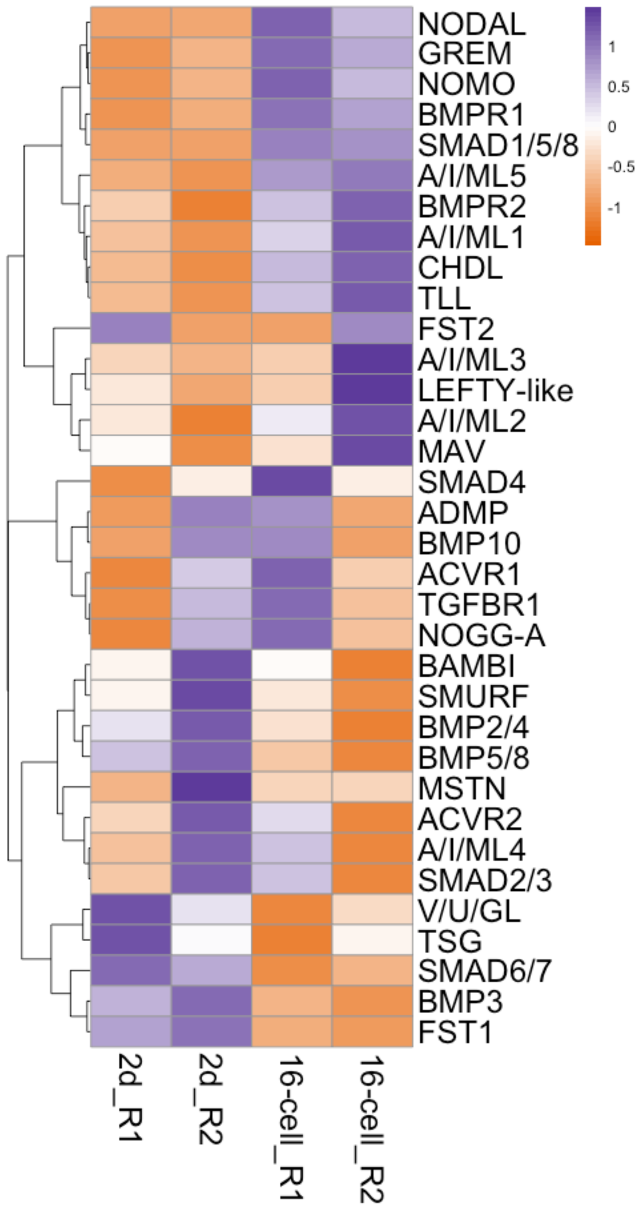


**Figure S4.**

Heatmap for z-scores of average normalized counts of TGF-ß components in all samples used for 2d vs 16-cell pairwise comparison. Black circles: differentially-enriched transcripts in 2d vs 16-cell.

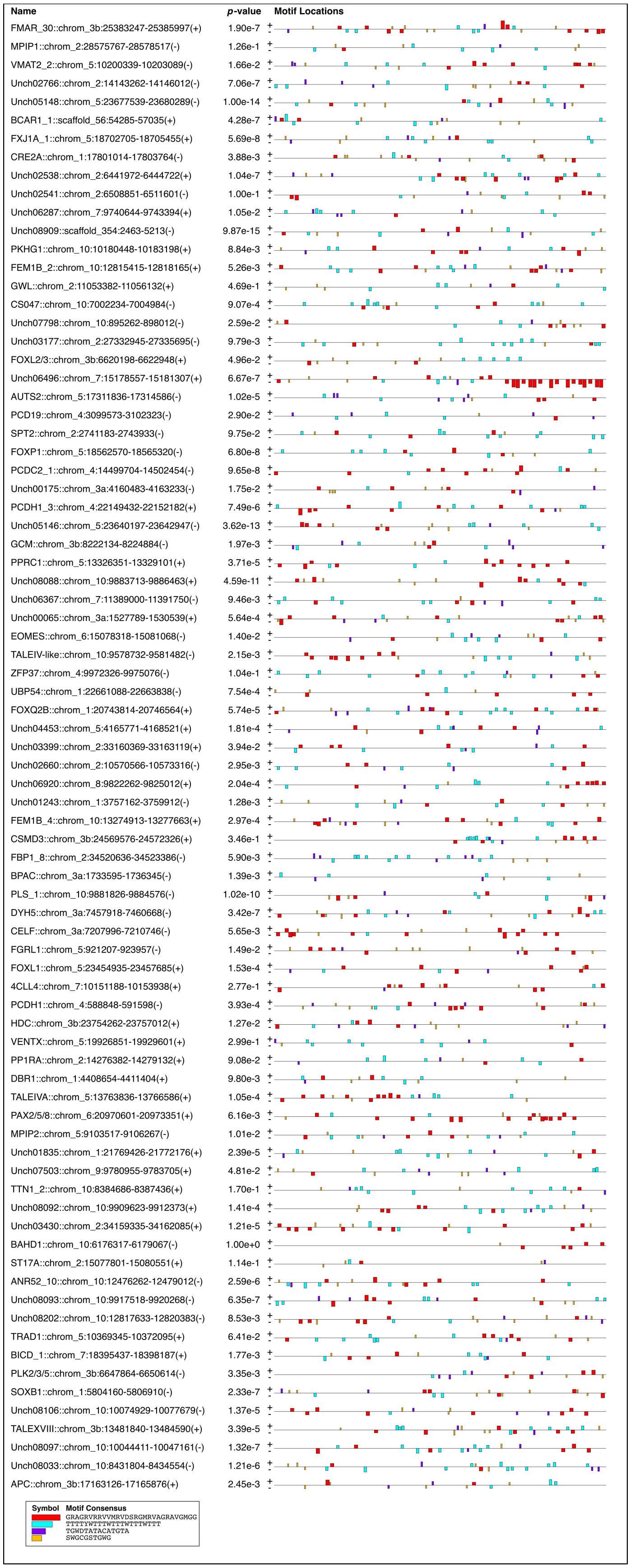
Name
p-value
Motif Locations
FMAR_30::chrom_3b:25383247-25385997(+) 1.90e-7
MPIP1::chrom_2:28575767-28578517(-) 1.26e-1
VMAT2_2::chrom_5:10200339-10203089(-) 1.66e-2
Unch02766::chrom_2:14143262-14146012(-) 7.06e-7
Unch05148::chrom_5:23677539-23680289(-) 1.00e-14
BCAR1_1::scaffold_56:54285-57035(+) 4.28e-7
FXJ1A_1::chrom_5:18702705-18705455(+) 5.69e-8
CRE2A::chrom_1:17801014-17803764(-) 3.88e-3
Unch02538::chrom_2:6441972-6444722(+) 1.04e-7
Unch02541::chrom_2:6508851-6511601(-) 1.00e-1
Unch06287::chrom_7:9740644-9743394(+) 1.05e-2
Unch08909::scaffold_354:2463-5213(-) 9.87e-15
PKHG1::chrom_10:10180448-10183198(+) 8.84e-3
FEM1B_2::chrom_10:12815415-12818165(+) 5.26e-3
GWL::chrom_2:11053382-11056132(+) 4.69e-1
CS047::chrom_10:7002234-7004984(-) 9.07e-4
Unch07798::chrom_10:895262-898012(-) 2.59e-2
Unch03177::chrom_2:27332945-27335695(-) 9.79e-3
FOXL2/3::chrom_3b:6620198-6622948(+) 4.96e-2
Unch06496::chrom_7:15178557-15181307(+) 6.67e-7
AUTS2::chrom_5:17311836-17314586(-) 1.02e-5
PCD19::chrom_4:3099573-3102323(-) 2.90e-2
SPT2::chrom_2:2741183-2743933(-) 9.75e-2
FOXP1::chrom_5:18562570-18565320(-) 6.80e-8
PCDC2_1::chrom_4:14499704-14502454(-) 9.65e-8
Unch00175::chrom_3a:4160483-4163233(-) 1.75e-2
PCDH1_3::chrom_4:22149432-22152182(+) 7.49e-6
Unch05146::chrom_5:23640197-23642947(-) 3.62e-13
GCM::chrom_3b:8222134-8224884(-) 1.97e-3
PPRC1::chrom_5:13326351-13329101(+) 3.71e-5
Unch08088::chrom_10:9883713-9886463(+) 4.59e-11
Unch06367::chrom_7:11389000-11391750(-) 9.46e-3
Unch00065::chrom_3a:1527789-1530539(+) 5.64e-4
EOMES::chrom_6:15078318-15081068(-) 1.40e-2
TALEIV-like::chrom_10:9578732-9581482(-) 2.15e-3
ZFP37::chrom_4:9972326-9975076(-) 1.04e-1
UBP54::chrom_1:22661088-22663838(-) 7.54e-4
FOXQ2B::chrom_1:20743814-20746564(+) 5.74e-5
Unch04453::chrom_5:4165771-4168521(+) 1.81e-4
Unch03399::chrom_2:33160369-33163119(+) 3.94e-2
Unch02660::chrom_2:10570566-10573316(-) 2.95e-3
Unch06920::chrom_8:9822262-9825012(+) 2.04e-4
Unch01243::chrom_1:3757162-3759912(-) 1.28e-3
FEM1B_4::chrom_10:13274913-13277663(+) 2.97e-4
CSMD3::chrom_3b:24569576-24572326(+) 3.46e-1
FBP1_8::chrom_2:34520636-34523386(-) 5.90e-3
BPAC::chrom_3a:1733595-1736345(-) 1.39e-3
PLS_1::chrom_10:9881826-9884576(-) 1.02e-10
DYH5::chrom_3a:7457918-7460668(-) 3.42e-7
CELF::chrom_3a:7207996-7210746(-) 5.65e-3
FGRL1::chrom_5:921207-923957(-) 1.49e-2
FOXL1::chrom_5:23454935-23457685(+) 1.53e-4
4CLL4::chrom_7:10151188-10153938(+) 2.77e-1
PCDH1::chrom_4:588848-591598(-) 3.93e-4
HDC::chrom_3b:23754262-23757012(+) 1.27e-2
VENTX::chrom_5:19926851-19929601(+) 2.99e-1
PP1RA::chrom_2:14276382-14279132(+) 9.08e-2
DBR1::chrom_1:4408654-4411404(+) 9.80e-3
TALEIVA::chrom_5:13763836-13766586(+) 1.05e-4
PAX2/5/8::chrom_6:20970601-20973351(+) 6.16e-3
MPIP2::chrom_5:9103517-9106267(-) 1.01e-2
Unch01835::chrom_1:21769426-21772176(+) 2.39e-5
Unch07503::chrom_9:9780955-9783705(+) 4.81e-2
TTN1_2::chrom_10:8384686-8387436(+) 1.70e-1
Unch08092::chrom_10:9909623-9912373(+) 1.41e-4
Unch03430::chrom_2:34159335-34162085(+) 1.21e-5
BAHD1::chrom_10:6176317-6179067(-) 1.00e+0
ST17A::chrom_2:15077801-15080551(+) 1.14e-1
ANR52_10::chrom_10:12476262-12479012(-) 2.59e-6
Unch08093::chrom_10:9917518-9920268(-) 6.35e-7
Unch08202::chrom_10:12817633-12820383(-) 8.53e-3
TRAD1::chrom_5:10369345-10372095(+) 6.41e-2
BICD_1::chrom_7:18395437-18398187(+) 1.77e-3
PLK2/3/5::chrom_3b:6647864-6650614(-) 3.35e-3
SOXB1::chrom_1:5804160-5806910(-) 2.33e-7
Unch08106::chrom_10:10074929-10077679(-) 1.37e-5
TALEXVIII::chrom_3b:13481840-13484590(+) 3.39e-5
Unch08097::chrom_10:10044411-10047161(-) 1.32e-7
Unch08033::chrom_10:8431804-8434554(-) 1.21e-6
APC::chrom_3b:17163126-17165876(+) 2.45e-3
Symbol
Motif Consensus
GRAGRVRRVVMRVDSRGMRVAGRAVGMGG
TTTTYWTTTWTTTWTTTWTTT
TGWDTATACATGTA
SWGCGSTGWG

**Figure S5.**

Motif enrichment analysis via MEME for -2ACD (vs. 16-cell) differentially-enriched transcripts. Column 1 represents the name of the gene, assigned based on their localization in the chromosome after extraction using bedtools. Column 2 is the p-value, representing the probability of a random motif in the sequence to match the motif. Column 3 shows the localization of the motifs. Four enriched motifs are displayed as squares with different colors, and visualized over the 2750 bp region to observe specific patterns of clustering across different genes.

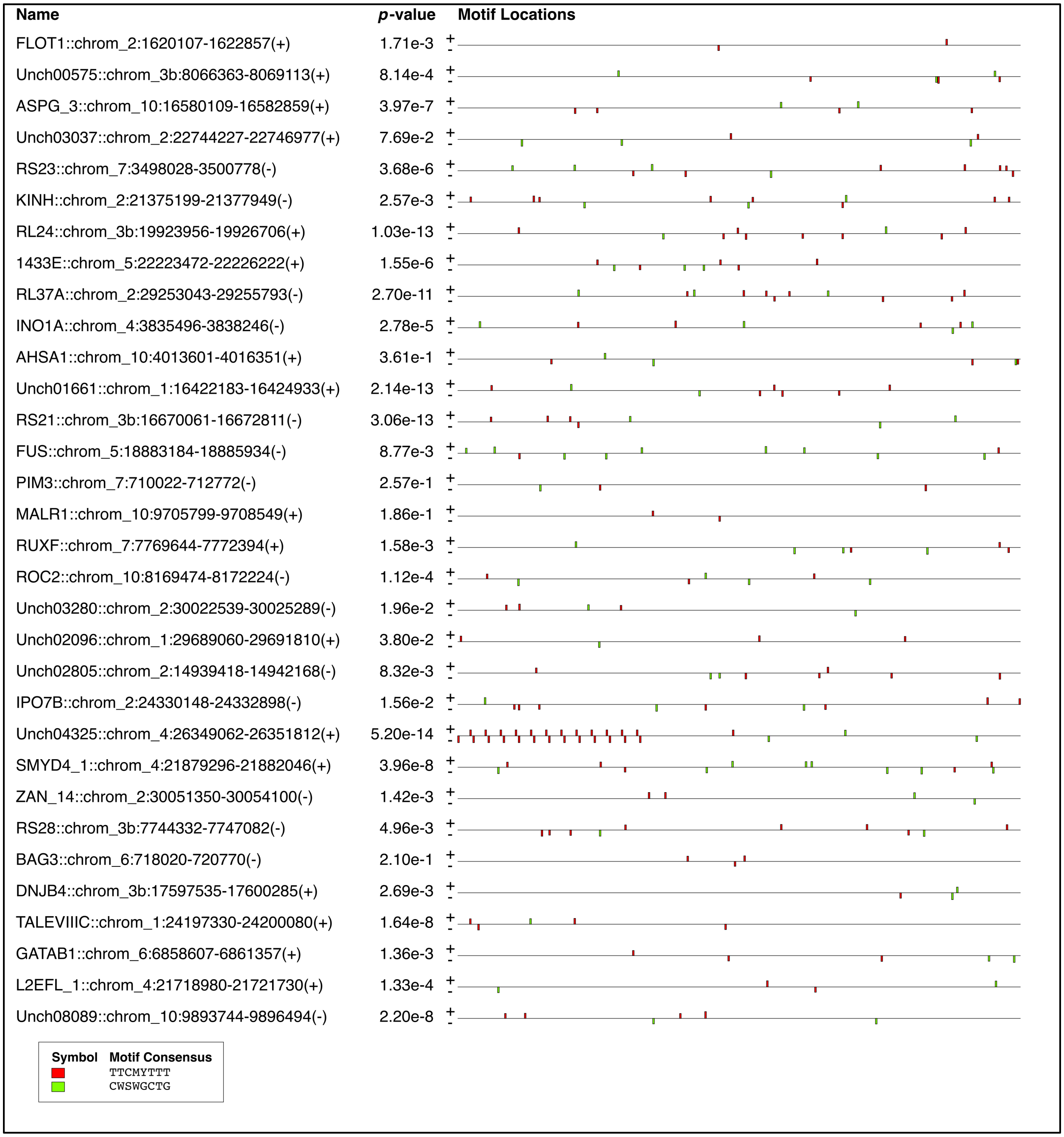
Name
p-value
Motif Locations
FLOT1::chrom_2:1620107-1622857(+)
1.71e-3
Unch00575::chrom_3b:8066363-8069113(+)
8.14e-4
ASPG_3::chrom_10:16580109-16582859(+)
3.97e-7
Unch03037::chrom_2:22744227-22746977(+)
7.69e-2
RS23::chrom_7:3498028-3500778(-)
3.68e-6
KINH::chrom_2:21375199-21377949(-)
2.57e-3
RL24::chrom_3b:19923956-19926706(+)
1.03e-13
1433E::chrom_5:22223472-22226222(+)
1.55e-6
RL37A::chrom_2:29253043-29255793(-)
2.70e-11
INO1A::chrom_4:3835496-3838246(-)
2.78e-5
AHSA1::chrom_10:4013601-4016351(+)
3.61e-1
Unch01661::chrom_1:16422183-16424933(+)
2.14e-13
RS21::chrom_3b:16670061-16672811(-)
3.06e-13
FUS::chrom_5:18883184-18885934(-)
8.77e-3
PIM3::chrom_7:710022-712772(-)
2.57e-1
MALR1::chrom_10:9705799-9708549(+)
1.86e-1
RUXF::chrom_7:7769644-7772394(+)
1.58e-3
ROC2::chrom_10:8169474-8172224(-)
1.12e-4
Unch03280::chrom_2:30022539-30025289(-)
1.96e-2
Unch02096::chrom_1:29689060-29691810(+)
3.80e-2
Unch02805::chrom_2:14939418-14942168(-)
8.32e-3
IPO7B::chrom_2:24330148-24332898(-)
1.56e-2
Unch04325::chrom_4:26349062-26351812(+)
5.20e-14
SMYD4_1::chrom_4:21879296-21882046(+)
3.96e-8
ZAN_14::chrom_2:30051350-30054100(-)
1.42e-3
RS28::chrom_3b:7744332-7747082(-)
4.96e-3
BAG3::chrom_6:718020-720770(-)
2.10e-1
DNJB4::chrom_3b:17597535-17600285(+)
2.69e-3
TALEVIIIC::chrom_1:24197330-24200080(+)
1.64e-8
GATAB1::chrom_6:6858607-6861357(+)
1.36e-3
L2EFL_1::chrom_4:21718980-21721730(+)
1.33e-4
Unch08089::chrom_10:9893744-9896494(-)
2.20e-8
Symbol
Motif Consensus
TTCMYTTT
CWSWGCTG

**Figure S6.**

Motif enrichment analysis via MEME for 16-cell (vs. -2ACD) differentially-enriched transcripts. Column 1 represents the name of the gene, assigned based on their localization in the chromosome after extraction using bedtools. Column 2 is the p-value, representing the probability of a random motif in the sequence to match the motif. Column 3 shows the localization of the motifs. Two enriched motifs are displayed as squares with different colors, and visualized over the 2750 bp region to observe specific patterns of clustering across different genes.

**Table S1.**

Comparison of nomenclature of isolations in Webster et al. (2026) and this article.

**Table S2.**

Differentially-expressed genes (DEGs) for all comparisons included in this article based on p-value $< 0.05$ and LFC $\geq 0.17$, with a thick line separating up- and down-regulated genes. Rows were organized in descending order based on log-fold change value. Symbol indicates the gene symbol from the top hit using BLASTp, identityVP indicates genes identified via orthology analysis, and PFAM indicates all domains found for each gene. Each sheet represents individual pairwise comparisons.

**Table S3.**

Motif occurrence search via FIMO, filtered for Mad/SMAD binding sites. Columns: (1) Motif ID, (2) Motif Alternative ID, (3) Gene, (4) Start of the motif in 2750 bp region, (5) End of the motif in the 2750 bp region, (6) Strand, (7) Score of the motif, (8) p-value, (9) q-value, (10) Matched sequence. Sheets: -1Q enriched transcripts, 8-cell enriched transcripts.

**Table S4.**

Species used for genomic analysis within Pleistoannelida (Errantia and Sedentaria) and data availability.

**Table S5.**

Experimental setup and quality control across multiple stages of analysis including replicates, number of embryos per treatment, RNA yield, RNA integrity number via TapeStation, number of paired reads after Illumina paired-end 150 bp sequencing, percentage of mapped reads via STAR, and differentially-enriched transcripts, transcription factor classes and signaling pathway components.

**Table S6.**

Sequencing quality for each sample based on pass filter clusters and number of reads with ≥30% of Q30 base-quality.
